# Targeted isolation of panels of diverse human protective broadly neutralizing antibodies against SARS-like viruses

**DOI:** 10.1101/2021.09.08.459480

**Authors:** Wan-ting He, Rami Musharrafieh, Ge Song, Katharina Dueker, Longping V. Tse, David R. Martinez, Alexandra Schäfer, Sean Callaghan, Peter Yong, Nathan Beutler, Jonathan L. Torres, Reid M. Volk, Panpan Zhou, Meng Yuan, Hejuni Liu, Fabio Anzanello, Tazio Capozzola, Mara Parren, Elijah Garcia, Stephen A. Rawlings, Davey M. Smith, Ian A. Wilson, Yana Safonova, Andrew B. Ward, Thomas F. Rogers, Ralph S. Baric, Lisa E. Gralinski, Dennis R. Burton, Raiees Andrabi

## Abstract

The emergence of current SARS-CoV-2 variants of concern (VOCs) and potential future spillovers of SARS-like coronaviruses into humans pose a major threat to human health and the global economy ^1–7^. Development of broadly effective coronavirus vaccines that can mitigate these threats is needed ^8, 9^. Notably, several recent studies have revealed that vaccination of recovered COVID-19 donors results in enhanced nAb responses compared to SARS-CoV-2 infection or vaccination alone ^10–13^. Here, we utilized a targeted donor selection strategy to isolate a large panel of broadly neutralizing antibodies (bnAbs) to sarbecoviruses from two such donors. Many of the bnAbs are remarkably effective in neutralization against sarbecoviruses that use ACE2 for viral entry and a substantial fraction also show notable binding to non-ACE2-using sarbecoviruses. The bnAbs are equally effective against most SARS-CoV-2 VOCs and many neutralize the Omicron variant. Neutralization breadth is achieved by bnAb binding to epitopes on a relatively conserved face of the receptor binding domain (RBD) as opposed to strain-specific nAbs to the receptor binding site that are commonly elicited in SARS-CoV-2 infection and vaccination ^14–18^. Consistent with targeting of conserved sites, select RBD bnAbs exhibited *in vivo* protective efficacy against diverse SARS-like coronaviruses in a prophylaxis challenge model. The generation of a large panel of potent bnAbs provides new opportunities and choices for next-generation antibody prophylactic and therapeutic applications and, importantly, provides a molecular basis for effective design of pan-sarbecovirus vaccines.

## Introduction

Relatively early in the COVID-19 pandemic, it appeared that SARS-CoV-2 was a virus that might be particularly amenable to control by vaccination. Many different vaccine modalities, most notably mRNA vaccination, showed spectacular success in phase 3 protection studies ^19, 20^. The success was attributed at least in part to the ability of the different modalities to induce robust neutralizing antibody (nAb) responses ^21–23^. However, as the virus has now infected hundreds of millions worldwide, variants have arisen (variants of concern, VOCs), some of which show notable resistance to neutralization by immunodominant nAb responses induced through infection and vaccination ^1, 3–5, 24^. Current vaccines are still apparently largely effective in preventing hospitalization and death caused by VOCs ^25, 26^. However, as vaccine-induced nAb responses naturally decline, breakthrough infections are on the increase and there are concerns that these may become more prevalent and perhaps more clinically serious, and that more pathogenic and resistant VOCs may appear. There are also concerns that emerging SARS-like viruses may seed new pandemics from spillover events ^2, 6^.

These concerns drive a search for nAbs and vaccines that are effective against a greater diversity of sarbecoviruses. Indeed, several individual broadly neutralizing antibodies (bnAbs) have now been generated either by direct isolation of bnAbs from convalescent donors or from antibody engineering of more strain-specific nAbs to generate breadth ^27–35^. Ideally, large panels of bnAbs would provide more options in the use of such Abs for prophylaxis and therapy ^36^. Importantly, a range of bnAbs would allow for better definition of the requirements for neutralization breadth and more rational effective design of appropriate immunogens ^37, 38^. A range of bnAbs has been crucial in germline targeting approaches to HIV vaccine design ^37–41^. Inspired by the demonstration ^10–13, 30, 42–48^ of the strong serum nAb responses in individuals who are infected with SARS-CoV-2 and then receive an mRNA vaccine, we isolated and characterized 40 bnAbs from two COVID-19 convalescent donors who were recently vaccinated, many of which combine excellent potency and breadth to sarbecoviruses. *In vivo* evaluation of select RBD bnAbs in a prophylaxis challenge model showed robust protection against diverse ACE2-utlizing sarbecoviruses.

## Results

### Donors for bnAb isolation

To identify donors for bnAb isolation, we first screened sera from 3 different groups for SARS-CoV-2 neutralization. The groups were: i) COVID-19 convalescent donors (n = 21); ii) spike-mRNA vaccinated (2X) donors (n = 10) and iii) COVID-19 convalescent donors (n = 15) who had subsequently been mRNA vaccinated (1X) (Fig. 1a, Extended Data Fig. 1). Consistent with earlier studies, we observed significantly higher levels of plasma nAbs in donors who were previously infected and then vaccinated (“recovered-vaccinated”) compared to donors who were only infected or only vaccinated (Fig. 1a, Extended Data Fig. 1). To examine the breadth of nAb responses across these 3 groups, we tested sera for neutralization against ACE2 receptor-utilizing sarbecoviruses (Pang17, SARS-CoV-1 and WIV1) and against SARS-CoV-2 VOCs (B.1.1.7 (Alpha), B.1.351 (Beta), P.1 (Gamma) and B.1.617.2 (Delta) and B.1.1.529 (Omicron)) (Fig. 1b-c, Extended Data Fig. 1). Sera from recovered-vaccinated donors showed greater breadth of neutralization and more effective neutralization of VOCs than sera from donors who were only previously infected or only vaccinated. Consistent with previous studies, neutralization efficacy of recovered-vaccinated sera against VOCs was similar to that against the ancestral strain of SARS-CoV-2 ^12, 44, 49, 50^ (Fig. 1c, Extended Data Fig. 1). Neutralization of SARS-CoV-1, whose spike is phylogenetically distinct (∼15% divergent at the amino acid level) from SARS-CoV-2 (Extended Data Fig. 1) ^2, 51^, was relatively low but was clearly above background for about half of the recovered-vaccinated donors (Extended Data Fig. 1). None of the convalescent-only or vaccinated-only donor sera could neutralize SARS-CoV-1, as also noted by us earlier ^52^. Of note, many of the existing SARS-CoV-2 cross-reactive or cross-neutralizing antibodies were isolated from SARS-CoV-1 convalescent donors ^27, 32, 53–56^ and only more recently from SARS-CoV-2 infected donors ^30, 31, 34, 35, 57^. BnAbs have also been isolated from SARS-CoV-2 S-protein vaccinated macaques that show serum cross-neutralizing activity against SARS-CoV-1 ^52, 58–60^. Hence, SARS-CoV-1/2 cross-neutralization appears to be a good indicator of the presence of pan-sarbecovirus activity and greater CoV neutralization breadth ^61^. Accordingly, for isolation of bnAbs in this study, we focused on two SARS-CoV-2 infection recovered-vaccinated donors (CC25 and CC84) with the most potent SARS-CoV-1 cross-neutralizing antibody titers.

**Fig. 1.**
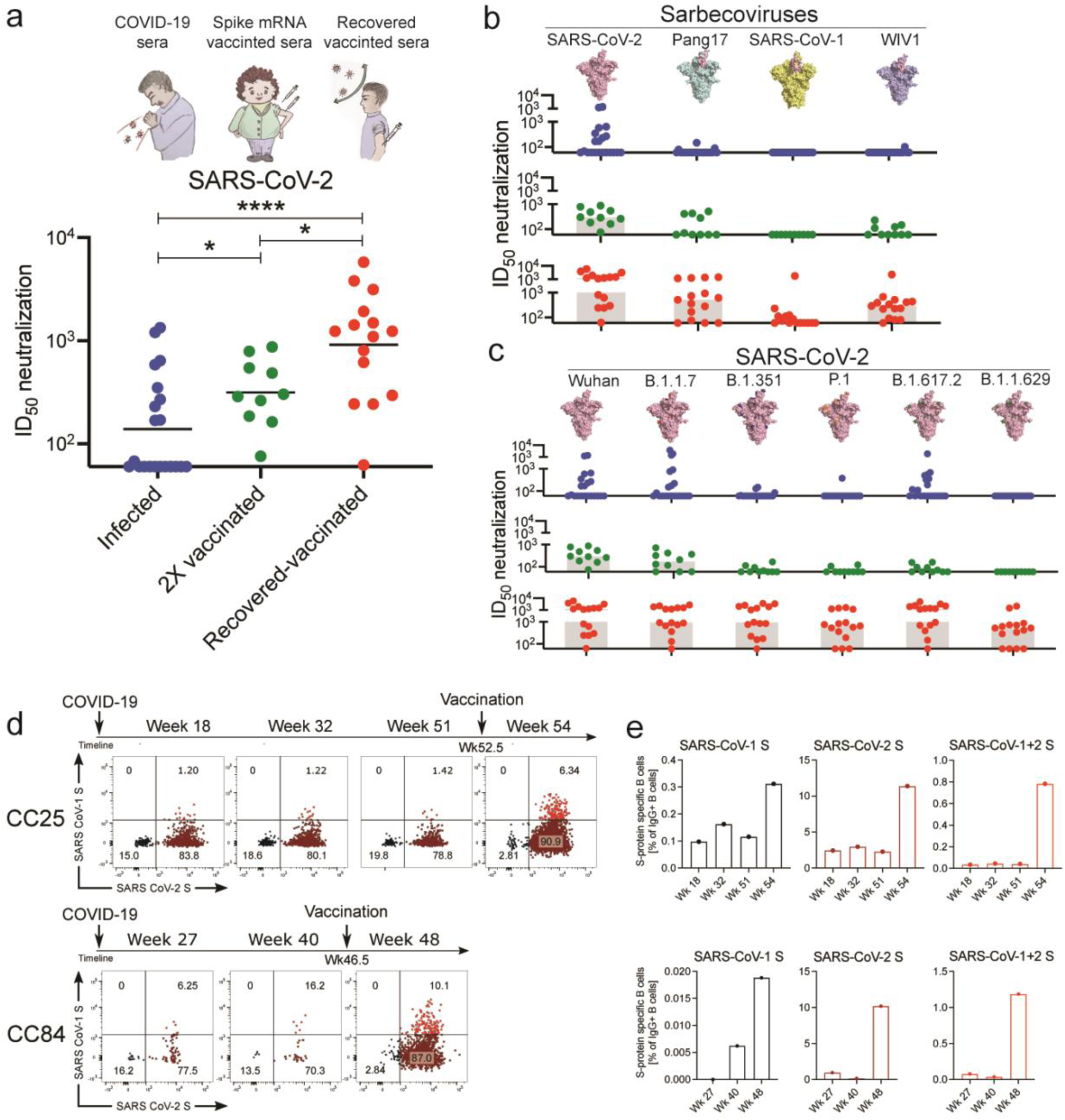
Plasma neutralization and B cell responses in convalescent recovered, vaccinated, and recovered-vaccinated donors. **a.** SARS-CoV-2 pseudovirus neutralization of plasma samples from COVID-19 convalescent recovered donors (blue: recovered), vaccinated donors (green: 2X vaccinated) or vaccinated donors with a prior history of SARS-CoV-2 infection (red: recovered-vaccinated). Statistical comparisons between the two groups were performed using a Mann-Whitney two-tailed test, (*p < 0.05, ****p < 0.0001). **b.** Plasma neutralization for all three groups against distantly related sarbecoviruses. Pang17 (cyan), SARS-CoV-1 (yellow), and WIV1 (violet) are shown. RBDs are colored pink for all spikes. In contrast to infected-only and vaccinated-only donors, approximately half of the recovered-vaccinated donors have neutralizing titers against SARS-CoV-1 above background (Extended Data Fig. 1). **c.** Plasma neutralizing activity against SARS-CoV-2 (Wuhan) and SARS-CoV-2 variants of concern (B.1.1.7 (Alpha), B.1.351 (Beta), P.1 (Gamma), B.1.617.2 (Delta) and B.1.1.529 (Omicron)). **d.** Flow cytometric analysis of IgG^+^ B cells from PBMCs of human donors CC25 and CC84 isolated at the indicated timepoints (see Extended Data Fig. 2 for gating strategy). Numbers indicate percentage of cells binding to SARS-CoV-1 and SARS-CoV-2 spike proteins, respectively. **e.** Quantification of SARS-CoV-1-specific, SARS-CoV-2-specific, and cross-reactive IgG^+^ B cells from donor CC25 (top) and CC84 (bottom) donors.

### Isolation and characterization of a large panel of sarbecovirus bnAbs

Using SARS-CoV-1 and SARS-CoV-2 S-proteins as baits, we sorted antigen-specific single B cells to isolate 107 mAbs from two COVID-19 recovered donors who had been recently vaccinated with the Moderna mRNA-1273 vaccine (CC25 (n = 56) and CC84 (n = 51)) (Extended Data Fig. 2) ^62^. Briefly, from the peripheral blood mononuclear cells (PBMCs) of the donors, we sorted CD19^+^CD20^+^ IgG^+^ IgM^-^ B cells that bound to both SARS-CoV-2 and SARS-CoV-1 S-proteins (Fig. 1d, Extended Data Fig. 2). Flow cytometry profiling of pre-vaccination (post-infection) PMBCs of CC25 and CC84 donors revealed that SARS-CoV-1/2 cross-reactive IgG^+^ B cells were likely seeded after infection and were recalled upon vaccination (Fig. 1d-e, Extended Data Fig. 2), as also observed in other studies ^44, 63^. Heavy (HC) and light (LC) chain sequences from 107 S-protein sorted single B cells were recovered and expressed as IgGs (Extended Data Fig. 3).

**Fig. 2.**
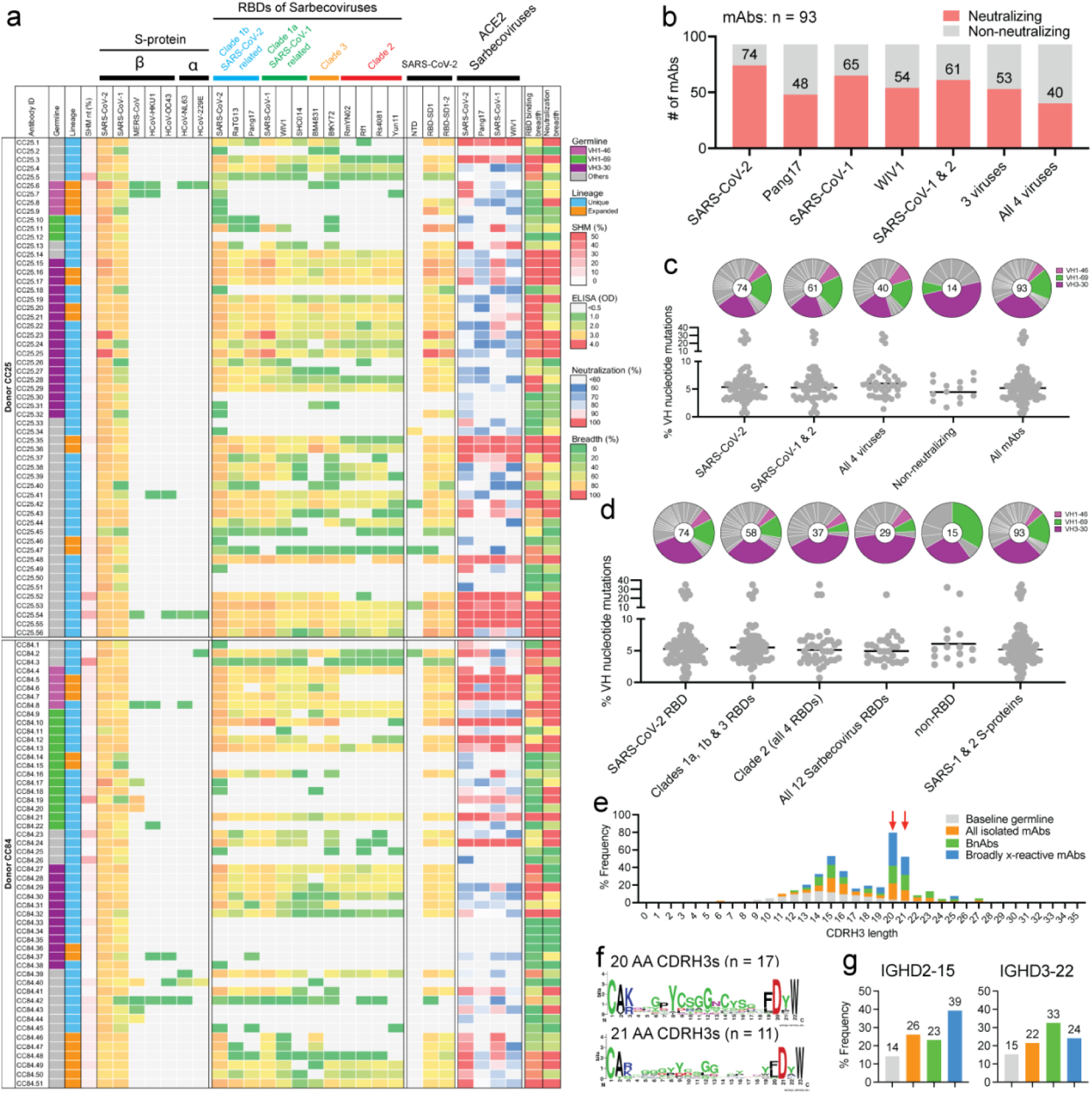
Binding, neutralization and immunogenetic properties of sarbecovirus mAbs. A total of 107 mAbs were isolated, 56 mAbs from donor CC25 and 51 mAbs from donor CC84. MAbs were isolated by single B cell sorting using SARS-CoV-1 and SARS-CoV-2 S-proteins as baits. **a.** Heatmap showing IGHV germline gene usage (colored: VH1-46 (magenta), VH1-69 (spring) and VH3-30 (plum) and other V-genes (grey)), lineage information (unique (sky) and expanded (tangerine) lineages) and V-gene nucleotide somatic mutations (SHMs). ELISA binding activity of mAbs with SARS-CoV-2, SARS-CoV-1 and other β- and α-HCoV derived S-proteins and domains of SARS-CoV-2 S-protein (NTD, RBD-SD1, RBD-SD1-2) (LOD <0.5 OD_405_). Binding of mAbs with clade 1a (SARS-CoV-2 related: SARS-CoV-2, RatG13, Pang17), clade 1b (SARS-CoV-1 related: SARS-CoV-1, WIV1, SHC014), clade 2 (RmYN02, Rf1, Rs4081, Yun11) and clade 3 (BM4831, BtKY72) sarbecovirus S-protein derived monomeric RBDs. Percent neutralization of ACE2-utilizing sarbecoviruses (SARS-CoV-2, Pang17, SARS-CoV-1 and WIV1) by mAb supernatants (cut-off <60%). Breadth (%) of binding to 12 sarbecovirus RBDs and breadth (%) of neutralization with 4 ACE2 sarbecoviruses is indicated for each mAb. MAb expression levels in the supernatants were quantified by total IgG ELISA and the concentrations were approximately comparable overall. For reproducibility, the binding and neutralization assays were all performed twice with mAb supernatants expressed independently twice. **b.** Number of mAbs (unique clones) neutralizing SARS-CoV-2 and other sarbecoviruses. 40 mAbs neutralized all 4 ACE2 sarbecoviruses tested. **c.** Pie plots showing IGHV gene usage distribution of isolated mAbs with enriched gene families colored, VH1-46 (magenta) VH1-69 (spring) and VH3-30 (plum). Dot plots showing % nucleotide mutations (SHMs) in the heavy chain (VH) of isolated mAbs. The mAbs are grouped by neutralization with sarbecoviruses. **d.** Pie and dot plots depicting IGHV gene distribution and VH nucleotide SHMs respectively. The mAbs are grouped by binding to sarbecovirus-derived RBDs. **e.** CDRH3 length distributions of isolated mAbs across broadly neutralizing and broadly cross-reactive mAb groups compared to human baseline germline reference. MAbs with 20- and 21- amino acid-CDRH3s are highly enriched. **f.** Sequence conservation logos of 20 (n = 17) and 21 (n = 11) amino acid long CDRH3-bearing mAbs show enrichment of D-gene derived residues, including IGHD2-15 germline D-gene encoded two cysteines in 20 amino acid long CDRH3-bearing mAbs that may potentially form a disulfide bond. **g.** Enrichment of IGHD2-15 and IGHD3-22 germline D-genes in sarbecovirus broadly neutralizing or broadly cross-reactive mAbs compared to corresponding human baseline germlines.

**Fig. 3.**
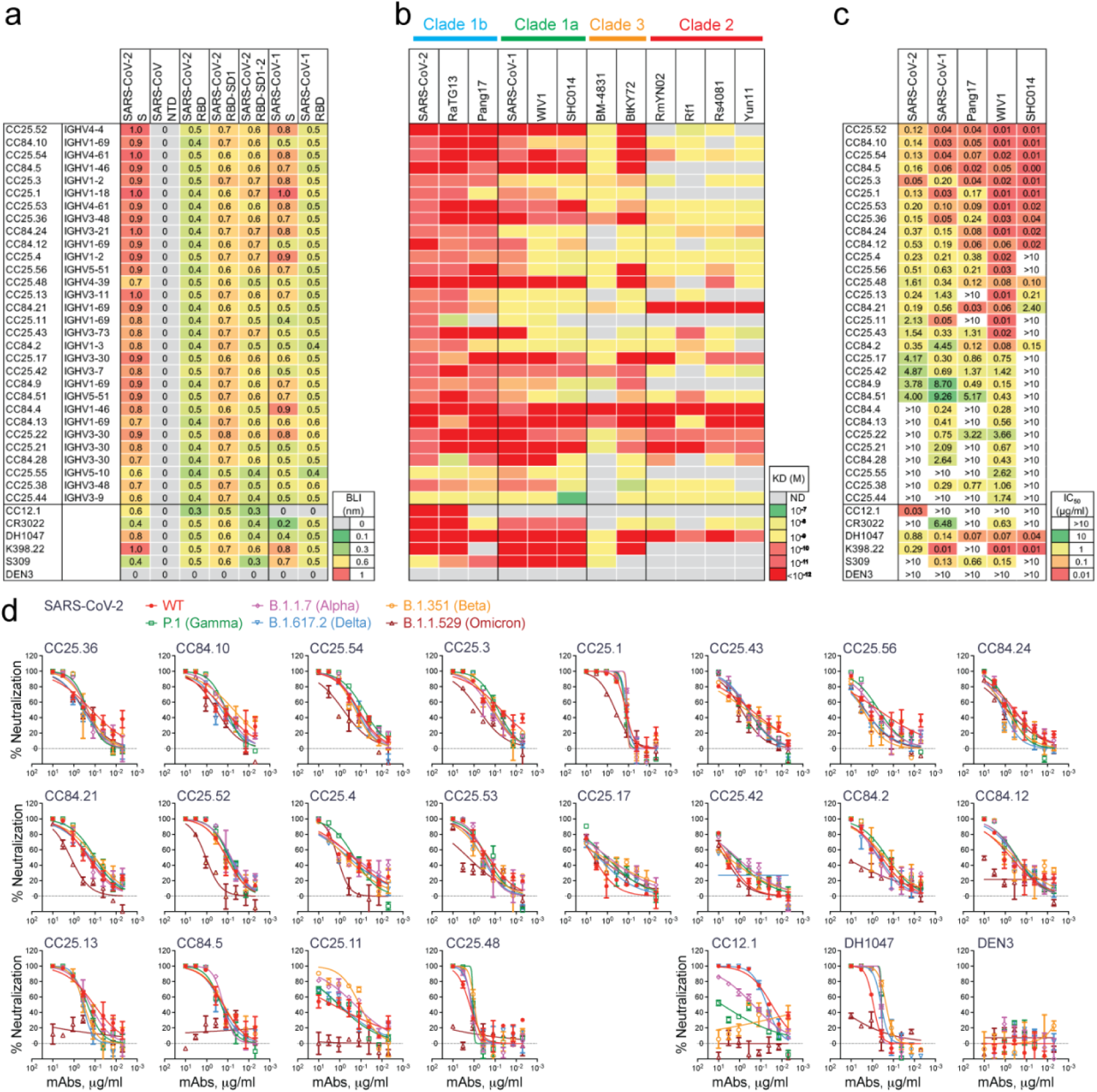
Binding and neutralization of mAbs in terms of affinity/potency and breadth. A total of 19 mAbs from donor CC25 and 11 mAbs from donor CC84 were selected to determine specificity, relative affinities and neutralization of sarbecoviruses and SARS-CoV-2 VOCs. **a.** Heatmap of binding responses (nm) determined by BLI using SARS-CoV-1 and SARS-CoV-2 S and S-protein domains and subdomains with IGHV gene usage for each mAb indicated. **b.** Heatmap of dissociation constants (K_D_ (M) values) for mAb binding to spike-derived monomeric RBDs from four sarbecovirus clades: clade 1b (n = 3); clade 1a (n = 3); clade 2 (n = 4); clade 3 (n = 2). Binding kinetics were obtained using the 1:1 binding kinetics fitting model on ForteBio Data Analysis software. **c.** IC_50_ neutralization of mAbs against SARS-CoV-2, SARS-CoV-1, Pang17, WIV1, and SHC014 determined using pseudotyped viruses. **d**, Neutralization of 20 bnAbs against SARS-CoV-2 (Wuhan) and five major SARS-CoV-2 variants of concern (B.1.1.7 (Alpha), B.1.351 (Beta), P.1 (Gamma), B.1.617.2 (Delta) and B.1.1.529 (Omicron)). SARS-CoV-2 Abs, CC12.1, DH1047, and Dengue Ab, DEN3 were used as controls.

All 107 mAbs exhibited cross-reactive ELISA binding to SARS-CoV-2 and SARS-CoV-1 S-proteins (Fig. 2a, Extended Data Fig. 3). Very few of the mAbs showed weak but detectable binding to β-HCoV (MERS-CoV HCoV-HKU1 and HCoV-OC43) and α-HCoV (HCoV-NL63 and HCoV-229E)-derived S-proteins (Fig. 2a, Extended Data Fig. 3). To determine the epitope specificities of the mAbs, we tested ELISA binding with SARS-CoV-2 S1 subunit domains and observed that the vast majority of the mAbs (>80%) displayed RBD-specific binding (Fig. 2a). To determine the cross-reactivity of RBD binding, we investigated 12 diverse RBDs representing all the 4 major sarbecovirus clades: clades 1a, 1b, 2 and 3 ^2, 27, 51^ (Fig. 2b, Extended Data Fig. 3). The mAbs showed the greatest degree of cross-reactivity with clades 1a and 1b and the least with clade 2. Approximately a third of the mAbs (31%) showed cross-reactivity with all 12 RBDs derived from all 4 sarbecovirus clades (Fig. 2b, Extended Data Fig. 3).

Next, we evaluated cross-clade neutralization with mAb supernatants on a panel of clade 1a (SARS-CoV-1 and WIV-1) and clade 1b (SARS-CoV-2 and Pang17) pseudoviruses of ACE2-utilizing sarbecoviruses ^2^. Two-thirds of mAbs neutralized both SARS-CoV-1 and SARS-CoV-2 and 43% (40 out of 93 mAbs) neutralized all 4 sarbecoviruses in the panel and are categorized as bnAbs in this study (Fig. 2b, Extended Data Fig. 3). More comprehensive and quantitative cross-neutralization is described with a smaller panel of mAbs below.

In terms of antibody sequences, of the 107 isolated mAbs, 93 were encoded by unique immunoglobulin germline gene combinations and 11 were expanded lineages (CC25 [n = 6] and CC84 [n = 5]) that had 2 or more clonal members (Fig. 2a, Extended Data Fig. 3). There was a notable enrichment of IGHV3-30, particularly, and also IGHV1-46 and IGHV1-69 germline gene families for both donors as compared to human baseline germline frequencies (Fig. 2a, c-d, Extended Data Fig. 4) ^64, 65^. Light chains of certain germline-gene families (IGKV1-33, IGKV2-30, IGLV1-40, IGLV3-21) were also modestly enriched in the isolated mAbs (Extended Data Fig. 4). Interestingly, the mAbs showed modest levels of V-gene nucleotide somatic hypermutation (SHM): for VH, median = 5.0% and for VL, median = 4.0% (Extended Data Fig. 3). As multiple studies have shown that heavy chains dominate the epitope interaction by RBD nAb ^5, 16, 17, 66^, we sought to determine whether the IGHV germline gene usage and/or VH SHM levels were correlated with the extent of neutralization breadth (Fig. 2a, c-d). We observed enrichment of IGHV3-30 in mAbs that bind to clade 2 RBDs and all 12 sarbecovirus RBDs, but otherwise no notable trends (Fig. 2c-d). VH-gene SHM levels did not distinguish potent broadly neutralizing from less broad or non-neutralizing mAbs. Overall, we observed that some IGHV or IGLV genes were enriched in bnAbs, and several human immunoglobulin gene combinations could encode for sarbecovirus bnAbs.

**Fig. 4.**
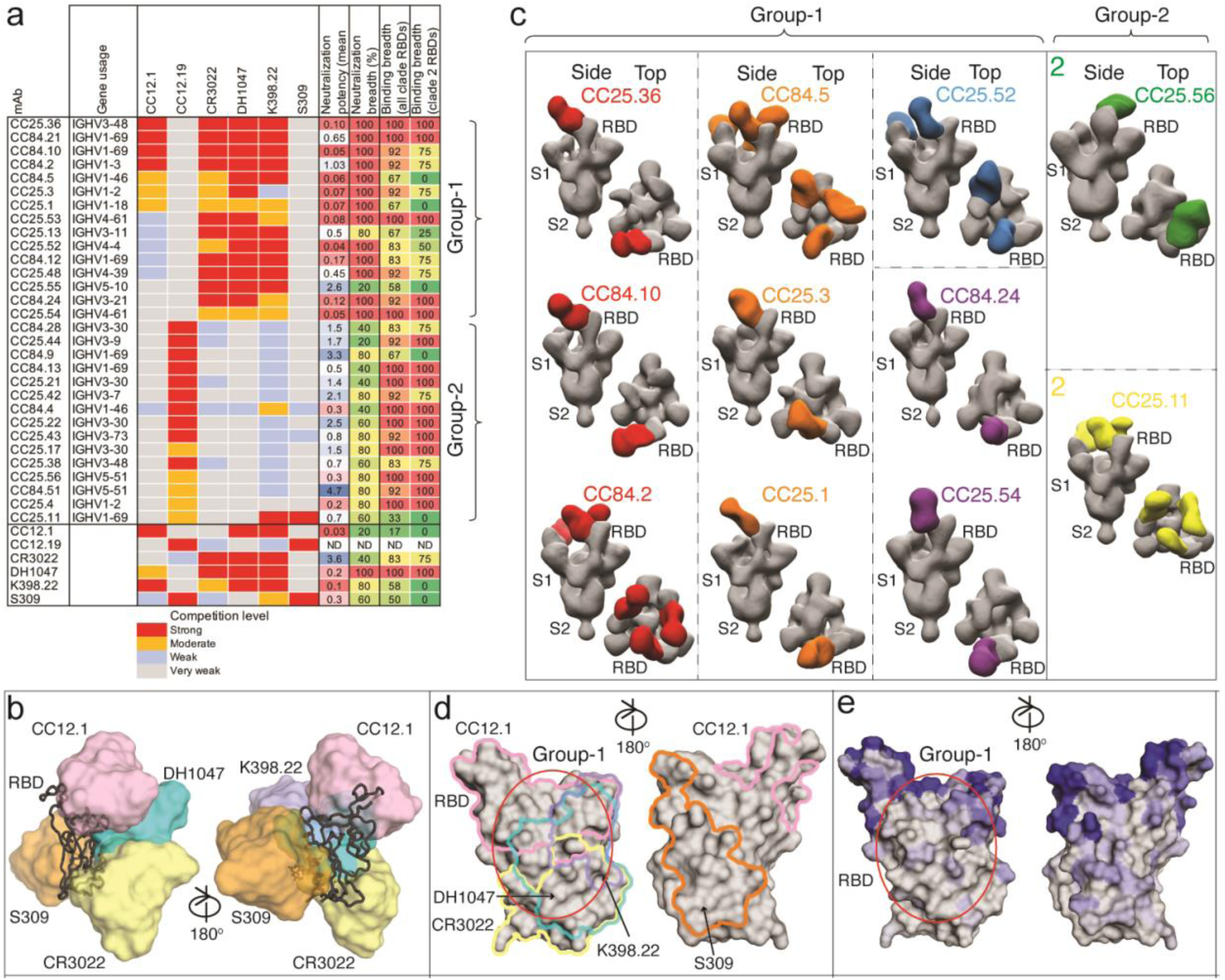
Epitope specificities of sarbecovirus bnAbs. **a.** Heatmap summary of epitope binning of sarbecovirus bnAbs based on BLI competition of bnAbs with human (CC12.1, CC12.19, CR3022, DH1047 and S309) and macaque (K398.22) RBD-specific nAbs. IGHV gene usage for each mAb is indicated. Geomean neutralization potency and breadth (calculated from Fig. 3c) and RBD binding breadth with clade 2 or all clade sarbecoviruses (calculated from Fig. 3b) for each mAb are indicated. The BLI competition was performed with monomeric SARS-CoV-2 RBD, and the competition levels are indicated as red (strong), orange (moderate), light blue (weak) and grey (very weak) competition. Based on competition with human and one macaque nAb of known specificities, the sarbecovirus bnAbs were divided into groups-1 and -2. **b.** Binding of human nAbs to SARS-CoV-2 RBD. The RBD is shown as a black chain trace, whereas antibodies are represented by solid surfaces in different colors: CC12.1 (pink, PDB 6XC2), CR3022 (yellow, PDB 6W41), S309 (orange, PDB 7R6W), DH1047 (cyan, 7LD1), and K398.22 (blue, PDB submitted) ^52^. **c.** Electron microscopy 3D reconstructions of sarbecovirus bnAb Fabs with SARS-CoV-2 S-protein. Fabs of group-1 (n = 9) and group-2 (n = 2) were complexed with SARS-CoV-2 S-protein and 3D reconstructions were generated from 2D class averages. Fabs are shown in different colors and the spike S1 and S2 subunits (grey) and the RBD sites are labelled. **d.** The epitope of each antibody is outlined in different colors corresponding to panel b. Epitope residues are defined by buried surface area (BSA) > 0 Å^2^ as calculated by PDBePISA (https://www.ebi.ac.uk/msd-srv/prot_int/pistart.html). Putative epitope regions of group-1 bnAbs based on the competitive binding assay are indicated by red circles. **e.** Conservation of 12 sarbecovirus RBDs. Gray surface represents conserved regions, whereas blue represents variable regions. The conservation was calculated by ConSurf (https://consurf.tau.ac.il/). The putative epitope region targeted by group-1 bnAbs is circled as in panel d.

To further investigate the potential contribution of SHM to broad reactivity to sarbecoviruses, we tested the binding of mAbs to SARS-CoV-2 S-protein and to monomeric RBD by BLI (Extended Data Fig. 5). We found no association of SHM with S-protein binding and a weak correlation with binding to RBD. Consistent with this lack of correlation, we did not observe any correlation of SARS-CoV-2 RBD mAb binding with sarbecovirus neutralization breadth or binding breadth, although some modest correlation was observed for S-protein binding (Extended Data Fig. 5). These results suggest that critical antibody paratope features for sarbecovirus breadth when targeting the sites described below may be germline-encoded and limited affinity maturation is needed. While recent findings demonstrate that the accumulation of SHM may increase potency and breadth ^13, 67^, this may not be a requisite feature of sarbecovirus bnAbs, as noted by others ^68^. The predominant use of certain germline gene segments in bnAbs suggests that a germline-targeting approach ^39–41^ to pan-sarbecovirus vaccines may be rewarding and the relatively low levels of SHM are promising for successful vaccine deployment provided appropriate immunogens can be designed.

**Fig. 5.**
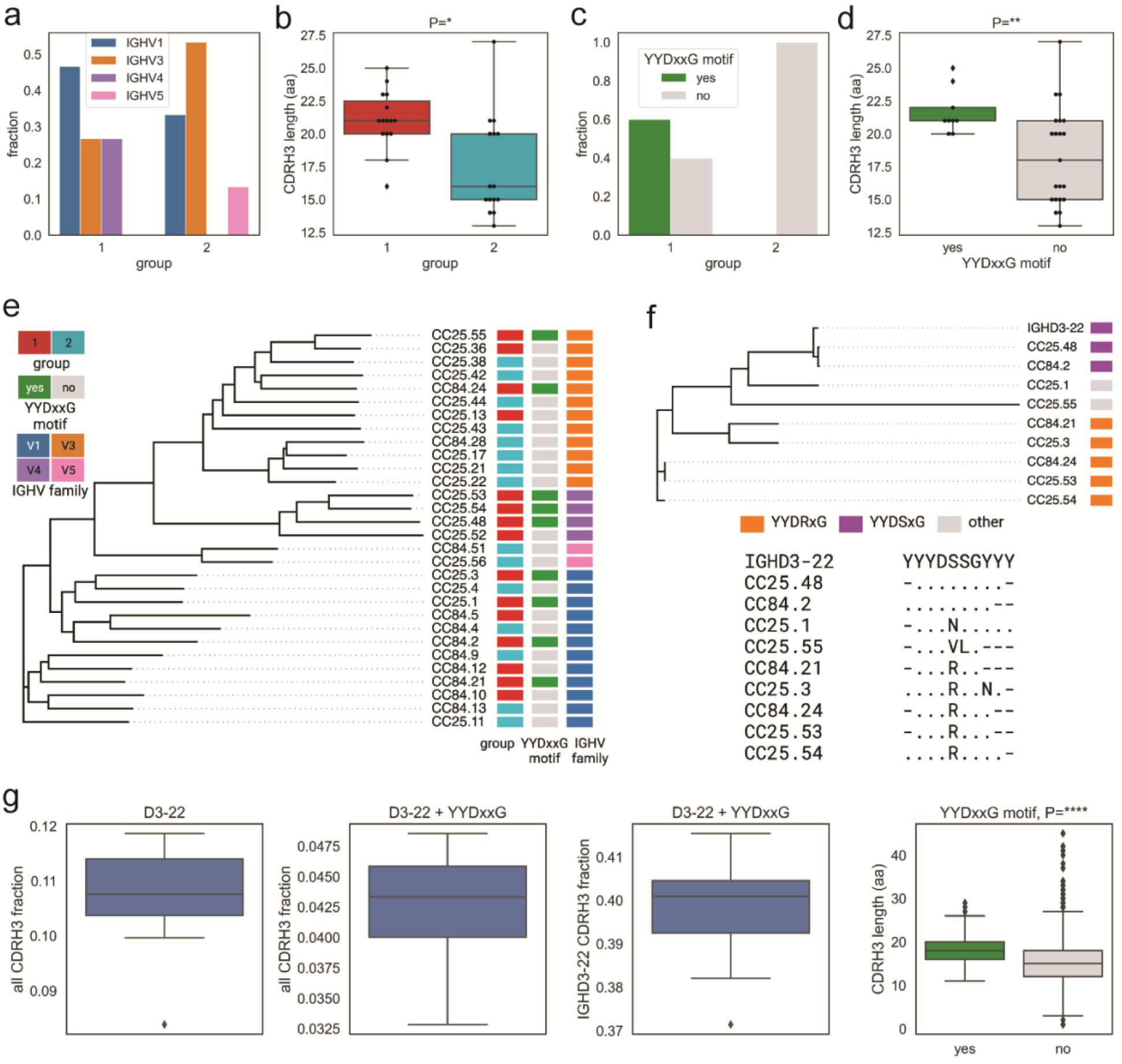
Immunogenetic properties of Group 1 and 2 RBD bnAbs. **a.** VH gene family usage: IGHV1 (blue), IGHV3 (orange), IGHV4 (violet), and IGHV5 (red). **b.** CDRH3 length distribution (amino acids) in Groups 1 (red) and 2 (cyan). P-values computed using the Kruskal-Wallis test and denoted as follows: P<0.05:*, P<0.005:**, P<0.00005:****. **c.** CDRH3 use of YYDxxG motif in group 1 and 2 RBD bnAbs: with (green) and without (gray). **d.** CDRH3 length distribution in RBD bnAbs with (green) and without (gray) YYDxxG motifs. **e.** Phylogenetic tree of heavy chain sequences of 30 RBD bnAbs. Each sequence is colored according to its group (left bar), IGHV gene family (middle bar), and the presence of the YYDxxD motif in the HCDR3 (right bar). Colors of heavy chain characteristics are consistent with panels **a-d**. Here and further, the phylogenetic tree is computed using Clustal Omega ^84^. **f.** Phylogenetic tree combining IGHD gene fragments of CDRH3s of nine mAbs with YYDxxG motifs and the amino acid translation of the germline sequence of IGHD3-22-containing YYGSSG. Each sequence is colored according to the amino acid following YYD: S (violet), R (orange), or others (gray). The alignment corresponding to the tree is shown below. Dots represent amino acids matching the germline amino acids. Germline amino acids truncated in CDRH3s are shown by dashes. **g.** Frequencies of IGHD3-22 germline genes with differing characteristics in naive heavy chain repertoires. From left to right: all IGHD3-22 in all CDRH3s; IGHD3-22 with YYDxxG motif in all CDRH3s; IGHD3-22 with YYDxxG motif in CDRH3s derived from IGHD3-22; distribution of lengths (in amino acids) of CDRH3s with (green) and without (gray) YYDxxG motifs. The fraction statistics were computed using ten Rep-seq libraries representing ten donors from the study by ^85^: ERR2567178–ERR2567187. The distribution of CDRH3 lengths was computed for library ERR2567178.

Next, we examined the CDRH3 loop lengths of the isolated Abs and observed a strong enrichment for 20- and 21-residue long CDRH3s compared to the human baseline reference database (Fig. 2e) ^64, 65^. These long CDRH3s were found to contain high proportions of two D genes, IGH D2-15 and D3-22, that were notably enriched in bnAbs (Fig. 2g). Notably, 71% (12/17) of mAbs with 20 amino acid CDRH3s utilized the germline IGHD2-15 D-gene, and the majority of mAbs bearing 21-amino acid-CDRH3s utilized either the IGHD2-15 or IGHD3-22 germline D-genes (Fig. 2f, Extended Data Figs. 3 and 4). We noted that the D3-22 D-gene was also selected in bnAbs isolated in other studies ^31, 57, 69, 70^. Therefore, vaccine design strategies will likely need to take these germline features into consideration ^39–41^.

Altogether, we have isolated a large panel of human sarbecovirus bnAbs. The isolated bnAbs, although encoded by several immunoglobulin germline gene families, are strongly enriched for certain germline gene features that will inform pan-sarbecovirus vaccine strategies.

### Detailed binding and neutralization characteristics of a smaller panel of bnAbs

We selected 30 SARS-CoV-1/SARS-CoV-2 RBD cross-reactive mAbs for more detailed characterization (Fig. 3a). Selection of mAbs was made based on a high degree of cross-reactive binding with RBDs of multiple sarbecovirus clades. The large panel above included nAbs that likely had more potent neutralization of SARS-CoV-1 and/or SARS-CoV-2 individually but lacked broad binding activity (Fig. 2a). To determine sarbecovirus binding cross-reactivity more extensively, we evaluated 12 soluble monomeric RBDs representing the major sarbecovirus clades, as above in Figure 2. Almost all mAbs bind SARS-CoV-2 and other clade 1b-derived RBDs, with most binding in a nanomolar (nM) to picomolar (pM) K_D_ affinity range (Fig. 3b, Extended Data Fig. 6). The mAbs that bound most effectively to clade 1b RBDs tended to also bind well to clade 1a and clade 3 RBDs, albeit with somewhat lower affinities, yet still in the nM-pM K_D_ affinity range. Cross-reactive binding was least to the clade 2 RBDs, although there was generally some level of reactivity and some mAbs did show high affinity binding to clade 2 RBDs. Remarkably, several mAbs showed consistently high affinity binding to RBDs from all 4 sarbecovirus clades (Fig. 3b, Extended Data Fig. 6).

**Fig. 6.**
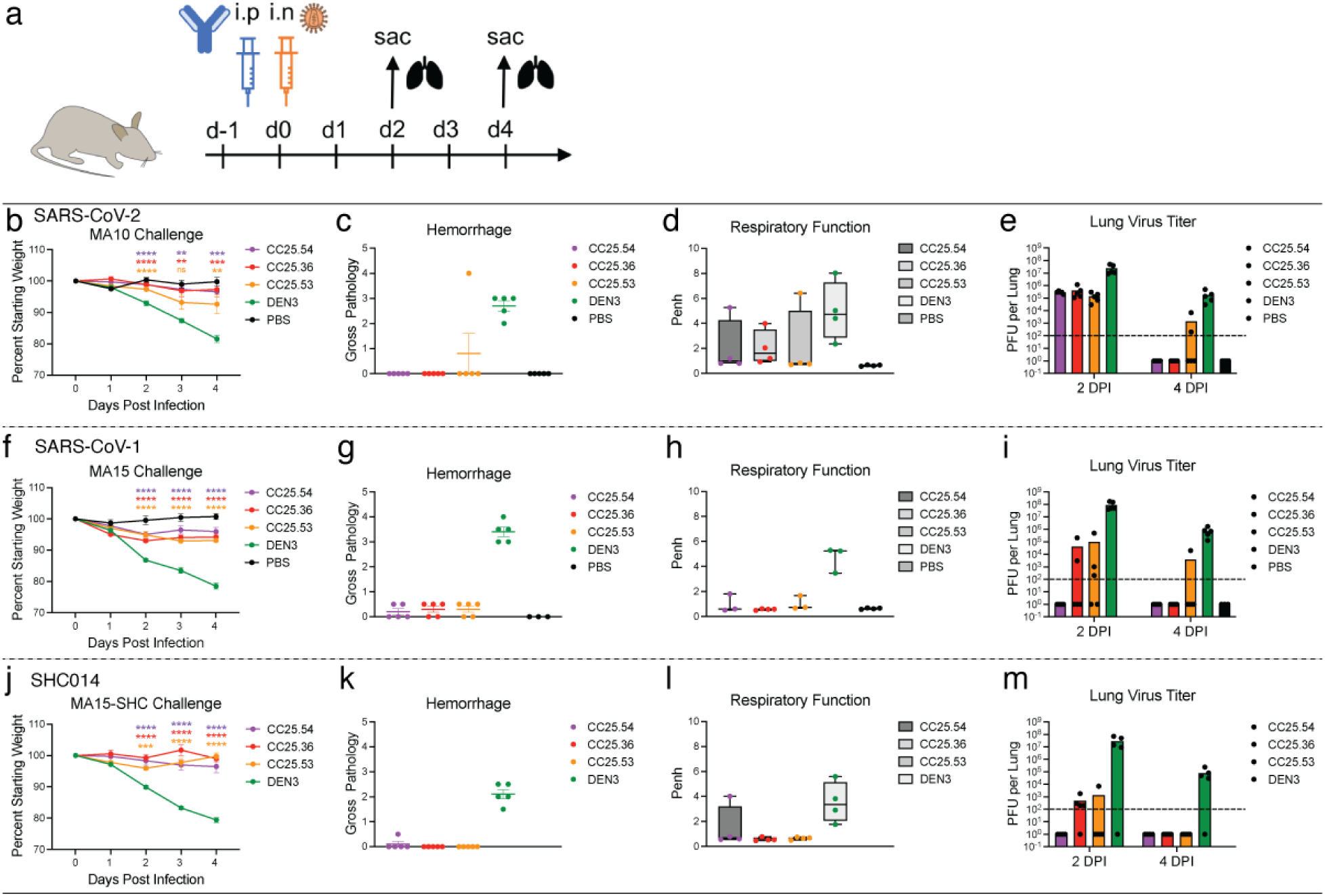
Prophylactic treatment of aged mice with RBD bnAbs protects against challenge with diverse SARS-like viruses. **a.** Three RBD bnAbs (CC25.54, CC25.36 and CC25.53) individually, or a dengue DEN3 control antibody were administered intra-peritoneally (i.p.) at 300μg per animal into 12 groups of aged mice (10 animals per group). Each group of animals was challenged intra-nasally (i.n.) 12h after antibody infusion with one of 3 mouse-adapted (MA) sarbecoviruses, (MA10 = SARS-CoV-2; 1 × 10^3^ plaque forming units (PFU), MA15 = SARS-CoV-1; 1 × 10^3^ PFU or MA15-SHC = SARS-CoV MA15-SHC014 chimera; 1 × 10^5^ PFU). As a control, groups of mice were exposed only to PBS in the absence of virus**. b,f,j.** Percent weight change in RBD bnAbs or DEN3 control antibody-treated animals after challenge with mouse-adapted sarbecoviruses. Percent weight change was calculated from day 0 starting weight for all animals. **c,g,k.** Day 2 post-infection Hemorrhage (Gross Pathology score) scored at tissue harvest in mice prophylactically treated with RBD bnAbs or DEN3 control mAb. **d,h,l.** Day 2 post-infection pulmonary function (shown as Penh score) was measured by whole body plethysmography in mice prophylactically treated with RBD bnAbs or DEN3 control mAb. **e,i,m.** Lung virus titers (PFU per Lung) were determined by plaque assay of lung tissues collected at day 2 or day 4 after infection. Statistical significance was calculated with Dunnett’s multiple comparisons test between each experimental group and the DEN3 control Ab group. (**p <0.01, ***p <0.001; ****p < 0.0001; ns-p >0.05).

Neutralization was investigated only for clade 1a and 1b ACE2-utilizing viruses, since neutralization assays were not available for clade 2 and 3 viruses. 22 of 30 mAbs neutralized SARS-CoV-2 with a range of IC_50_ neutralization titers (IC_50_ range = 0.05-4.9 µg/mL) (Fig. 3c) and 28 of 30 mAbs neutralized SARS-CoV-1, including all mAbs that neutralized SARS-CoV-2. Neutralization potency was typically stronger against SARS-CoV-1 than SARS-CoV-2. All mAbs showed neutralization against WIV1, while a majority exhibited cross-neutralization with Pang17, and to a lesser degree with SHC014. 13 out of 30 mAbs neutralized all 5 ACE2-utilizing sarbecoviruses tested with a geomean IC_50_ potency of 0.12 µg/ml. The three most potent SARS-CoV-2 bnAbs, CC25.52, CC25.54 and CC25.3, neutralized all 5 ACE2-utilizing sarbecoviruses with geomean potencies of 0.03, 0.04 and 0.04 µg/ml, respectively. Although neutralization assays differ, this suggests they are amongst the most potent and broad individual nAbs described to date (compare also control nAbs in Fig. 3c).

We tested neutralization of SARS-CoV-2 VOCs by 20 select SARS-CoV-2 bnAbs. Consistent with the donor CC25 and CC84 sera neutralization above, the bnAbs were consistently effective against SARS-CoV-2 VOCs tested (Fig. 3d, Extended Data Fig. 7). The IC_50_ neutralization titers of bnAbs remained largely unchanged against the Alpha, Beta, Gamma and Delta SARS-CoV-2 VOCs but were more affected by substitutions in the Omicron variant (Fig. 3d, Extended Data Fig. 7). Nevertheless, IC_50_ neutralization titers for many bnAbs were unchanged or minimally affected for the Omicron variant and remarkably 14 of 20 bnAbs retained significant neutralization against this highly evolved SARS-CoV-2 variant (Fig. 3d, Extended Data Fig. 7). In comparison, SARS-CoV-2 strain-specific nAb, CC12.1 showed substantial or complete loss of neutralization with VOCs. The results suggest that these bnAbs target more conserved RBD epitopes that are likely more resistant to SARS-CoV-2 escape mutations. Overall, we have identified multiple potent sarbecoviruses bnAbs that exhibit broad reactivity to SARS-CoV-2 variants and diverse sarbecovirus lineages.

### Epitope specificity of sarbecovirus bnAbs

To help map the epitopes recognized by the sarbecovirus bnAbs, we first epitope binned them using BLI competition with RBD nAbs of known specificities (Fig. 4a, Extended Data Fig. 8), including 5 human nAbs: (1) CC12.1, an RBS-A or class 1 nAb targeting the ACE2 binding site ^5, 14, 17^; (2) CC12.19, which is thought to recognize a complex RBD epitope and competes with some non-RBD Abs ^15^; (3) CR3022, which recognizes the class 4 epitope site ^5, 14^; (4) S309, which recognizes the class 3 epitope site ^5, 14^; and (5) DH1047, which recognizes a conserved site and is class 4 ^32^. In addition, we included K398.22, a macaque bnAb ^52^, which targets an RBD bnAb epitope distinct from that recognized by human bnAbs characterized to date but has features characteristic of class 4 bnAbs (Fig. 4a-b). The bnAbs we describe here can be clustered for convenience into two major groups. Group-1 bnAbs strongly competed with SARS-CoV-2 class 4 human bnAbs, CR3022 and DH1047, and macaque bnAb K398.22, showed more sporadic competition with CC12.1 and did not compete with CC12.19 or S309. Group-2 mAbs competed strongly with CC12.19, weakly with macaque K398.22, and only infrequently and/or weakly with any of the other bnAbs. Group-1 bnAbs were potent and broad in neutralization against ACE2-utilizing sarbecoviruses, but many lineage members displayed limited binding reactivity with clade 2 sarbecovirus RBDs. The group-2 mAbs showed broader binding reactivity with sarbecoviruses but were relatively less potent compared to group-1 bnAbs (Fig. 4a). Notably, one group-2 bnAb, CC25.11 showed strong competition with human class 3 RBD bnAb, S309 ^53^, and the macaque bnAb, K398.22 ^52^. The findings suggest that both group-1 and 2 bnAbs target more conserved RBD epitopes but group-1 bnAbs are overall more potent but less broad against clade 2 sarbecoviruses, with some exceptions.

To further investigate the epitopes recognized by the bnAbs, we utilized single-particle, negative-stain electron microscopy (nsEM) and confirmed that the 9 Group-1 and 2 Group-2 bnAb Fabs bound to the RBD of SARS-CoV-2 S-protein (Fig. 4c, Extended Data Fig. 9). The binding modes of bnAbs to SARS-CoV-2 S-protein were largely similar with some differences in the angles of approaches, but not distinct enough to clearly segregate group-1 epitope bnAbs. Further, structural studies that reveal molecular details of the antibody-antigen interactions contributing to the differences in the epitope recognition are important. The group-2 bnAb reconstructions are consistent with an epitope that spans the RBD, and other parts of the S-protein as described for the competitive Ab CC12.19^15^. These bnAb Fabs showed binding to S-protein with all three stoichiometries (Fab: trimer; 1:1, 1:2 and 1:3) with some of the Fabs exhibiting destabilizing effects on the S-trimer (Fig. 4c, Extended Data Fig. 9). This destabilization is seen as dimers and flexible densities in the 2D class averages (Extended Data Fig. 9).

### Immunogenetics of group 1 and 2 RBD bnAbs and vaccine targeting

To further understand the differences between group 1 and 2 RBD bnAbs and to determine if germline gene features can differentiate their epitope properties, we performed detailed antibody immunogenetic analysis. Both groups of RBD bnAbs were encoded by a number of IGHV germline gene families (Fig. 5a). The average CDRH3 loop lengths were significantly longer (p < 0.05) in group 1 compared to group 2 RBD bnAbs (Fig. 5b). Notably, group 1 bnAbs strongly enriched (60%: 9 out of 15 group 1 RBD bnAbs) for IGHD3-22 germline D-gene-encoded CDRH3 “YYDxxG” motifs and possessed significantly longer CDRH3 loops (p < 0.005) compared to the other mAbs (Fig. 5c-d). The IGHD3-22 germline D-gene “YYDxxG” motif-bearing group 1 RBD bnAbs utilized several IGHV germline gene combinations and the D-gene motifs were either retained in a germline configuration (YYDSSG: CC25.48 and CC84.2) or one or both “x” residues were mutated (Fig. 5c-d). The most common mutation was the substitution of YYD-proceeding x-residue, Serine-(S) to an Arginine-(R), which recurrently appeared in multiple YYDxxG motif bearing RBD bnAbs from both donors suggesting common B cell affinity maturation pathways. Interestingly, the S-R somatic mutation in the CDRH3 YYDxxG motif appeared to be important for resisting Omicron neutralization escape, as the non-mutated YYDxxG motif bearing group 1 RBD bnAbs failed or weakly neutralized this variant (Fig. 5d, Extended Data Fig. 7). Consistent with this observation, recent studies provide evidence for how YYDRxG RBD bnAbs can effectively bind to the conserved face of SARS-CoV-2 spike RBD ^71, 72^. The findings reveal that RBD bnAbs with certain recurrent germline features can effectively resist SARS-CoV-2 Omicron escape (Extended Data Fig. 7), but several antibody solutions can counter this extreme antigenic shift and should be considered for vaccine targeting. Remarkably, in our study genetically diverse RBD bnAbs in both groups 1 and 2 were capable of effectively neutralizing the Omicron variant (Extended Data Fig. 7) suggesting that several human antibody solutions can counter SARS-CoV-2 antigenic shift.

The association of a germline D-gene encoded motif, provides an opportunity for broad vaccine targeting, as has been described for HIV ^39, 41, 73^. For SARS-like coronaviruses, the YYDxxG motif ^71, 72^ appears promising for vaccine targeting. Encouragingly, human naïve B cell repertoires encode a sizable fraction of IGHD3-22 germline D-gene-encoded YYDxxG motif bearing B cells with desired CDRH3 lengths (Fig. 5g) that could be targeted by rationally designed vaccines ^74^.

### RBD bnAbs protect against challenge with diverse sarbecoviruses

To determine the protective efficacy of the RBD bnAbs, we conducted passive antibody transfer followed by challenge with sarbecoviruses in aged mice ^75^. We selected 3 of the broadest group-1 bnAbs, CC25.36, CC25.53 and CC25.54 and investigated their *in vivo* protective efficacy against SARS-CoV-2, SARS-CoV-1 and SHC014 sarbecoviruses in mice. SHC014 was chosen as it encodes extensive heterogeneity in the spike RBD, reduces mRNA SARS-CoV-2 polyclonal neutralization sera titers by ∼300-fold ^76^ and replicates efficiently in mice ^77^. Prior to the protection studies, we compared neutralization by RBD bnAbs of replication-competent viruses with that of pseudoviruses (Extended Data Fig. S10). Neutralization of replication-competent SARS-CoV-1 and SARS-CoV-2 by the bnAbs was more effective (lower IC_50_ values) than the corresponding pseudoviruses. Neutralization of replication-competent and pseudovirus versions of SHC014 by the bnAbs was approximately equivalent. The 3 RBD bnAbs, individually, or a DEN3 control antibody were administered intra-peritoneally (i.p.) at 300μg/animal into 12 groups of 10 animals (3 groups per antibody; Fig. 6a). Each group was challenged with one of 3 mouse-adapted (MA) sarbecoviruses, (MA10 = SARS-CoV-2, MA15 = SARS-CoV or MA15-SHC = SARS-CoV MA15 - SHC014 chimera), by intranasal (i.n.) administration of virus 12h post-antibody infusion (Fig. 6a). The animals were monitored for signs of clinical disease due to infection, including daily weight changes, and pulmonary function. Animals in each group were euthanized at day 2 or day 4 post infection and lung tissues were collected to determine virus titers by plaque assay. Gross pathology was also assessed at the time of tissue harvest. The RBD bnAb-treated animals in all 3 sarbecoviruses challenge experiments showed significantly reduced weight loss (Fig. 6b, f, j), reduced hemorrhage (Fig. 6c, g, k), and largely unaffected pulmonary function (Fig. 6d, h, l), as compared to the DEN3-treated control group animals, suggesting a protective role for bnAbs. We also examined virus load in the lungs at day 2 and day 4 post infection and, consistent with the above results, both the day 2 and day 4 viral titers in RBD bnAb-treated animals were substantially reduced compared to the DEN3-treated control group animals (Fig. 6e, i, m). Overall, all 3 RBD bnAbs protected against severe sarbecoviruses disease, CC25.54 and CC25.36 bnAbs being relatively more protective than CC25.53 bnAb. The animal data suggest potential utilization of the bnAbs in intervention strategies against diverse sarbecoviruses.

## Discussion

Here, we characterized a large panel of sarbecovirus bnAbs isolated from two SARS-CoV-2 recovered-vaccinated donors. Select bnAbs showed robust *in vivo* protection against diverse SARS-like viruses, including SARS-CoV-1, SARS-CoV-2 and SHC014, in a prophylaxis challenge model. The bnAbs are potent and show neutralization of a range of VOCs, and many are effective against Omicron. The bnAbs recognize a relatively conserved face of the RBD that overlaps with the footprint of a number of antibodies including ADG 61123, DH1047 and CR3022 ^32, 54, 72^ and broadly the face designated as that recognized by class 4 antibodies ^5, 66^. However, as illustrated in Fig 4, the panel of bnAbs differ in many details of recognition, for instance some compete with ACE2, others do not, and these differences are important in resistance to mutations. As variants such as Omicron emerge during this and future CoV pandemics, the availability of a selection of potent bnAbs provides choice of optimal reagents for antibody-based interventions to respond to the viral threats.

In terms of vaccine design, the generation of HIV immunogens typically draws heavily on the availability of multiple bnAbs to a given site to provide the best input for design strategies ^37, 78^. The same consideration is likely to apply to pan-sarbecovirus vaccine design. Further, although the bnAbs that we isolated were encoded by several gene families, certain V and D gene families were highly enriched. We confirmed and identified specific antibody germline gene features associated with broad activity against diverse sarbecoviruses and vaccine design strategies may seek to target these genetic features by rationally designed prophylactic vaccines ^37, 39–41, 74^. Some of the most potent bnAbs compete with the immunodominant human SARS-CoV-2 RBS-A/class 1 nAb CC12.1 that shows relatively low cross-reactivity. Elicitation of nAbs like CC12.1 may then reduce the elicitation of bnAbs, and rational vaccine design modalities may need to mask RBS-A/class 1 immunodominant sites ^79–81^ whilst leaving the bnAb sites intact. Resurfaced RBD-based immunogens in various flavors ^51, 58–60^ may achieve a similar goal.

Given the strong bnAb responses induced through infection-vaccination as indicated from serum studies and by our mAbs, are there lessons here for vaccine design? The higher frequency of bnAbs in infection-vaccination may have a number of causes. First, the spike S-protein may have subtle conformational differences, particularly in the sites targeted by bnAbs, between the native structure on virions and the stabilized form presented by mRNA immunization. This may favor the activation of bnAbs in the infection step followed by recall during mRNA boosting. Second, the long time-lag between infection and vaccination may have favored the accumulation of key mutations associated with bnAbs. There is evidence that intact HIV virions can be maintained on follicular dendritic cells in germinal centers over long time-periods in a mouse model ^82^. Third, T cell help provided by the infection may be superior to that provided by mRNA vaccination alone. Overall, there is an intriguing possibility that pan-sarbecovirus nAb activity may be best achieved by a hybrid approach ^43^ to immunization that seeks to mimic infection-vaccination, once the key contributing factors to breadth development in that approach can be determined. However, we also note that a very recent report, published as a resubmission of this manuscript was prepared, describes bnAbs arising from a third immunization with an inactivated vaccine ^83^.

In summary, we isolated multiple potent sarbecovirus protective cross-neutralizing human antibodies and provide a molecular basis for broad neutralization. The bnAbs identified may themselves have prophylactic utility and the bnAb panel delineates the boundaries and requirements for broad neutralization and will be an important contributor to rational vaccine design.

## Acknowledgements

We thank all the human cohort participants for donating samples. This work was supported by NIH CHAVD UM1 AI44462 (D.R.B.), NIH R61 AI161818 (R.A.), the IAVI Neutralizing Antibody Center, the Bill and Melinda Gates Foundation INV-004923 (I.A.W., A.B.W., D.R.B.), the Translational Virology Core of the San Diego Center for AIDS Research (CFAR) grant NIH AI036214 (D.M.S.), NIH 5T32AI007384 (S.A.R.), NIH AI149644 and AI157155 (R.S.B), NIH R21 AI145372 (L.E.G.), and the John and Mary Tu Foundation and the James B. Pendleton Charitable Trust (D.M.S. and D.R.B.). L.V.T. is supported by Pfizer NCBiotech Distinguished Postdoctoral Fellowship in Gene Therapy. D.R.M. is currently supported by a Burroughs Wellcome Fund Postdoctoral Enrichment Program Award and a Hanna H. Gray Fellowship from the Howard Hugues Medical Institute.

## Author contributions

W.H., R.M., G.S., K.D., D.R.B. and R.A. conceived and designed the study. N.B., M.P., E.G., S.A.R., D.M.S., and T.F.R. recruited donors and collected and processed plasma and PBMC samples. W.H., R.M., G.S., K.D., S.C., P.Y. and F.A. performed BLI, ELISA, virus preparation, neutralization and isolation and characterization of monoclonal antibodies. Y.S. performed immunogenetic analysis of antibodies. P.Z. prepared virus mutant plasmids. J.L.T. and R.M.V. conducted negative stain electron microscopy studies. M.Y. and H.L. generated antibody-antigen structural models. L.V.T. performed live virus neutralizations assays and L.V.T., D.R.M., A.S., and L.E.G. conducted *in vivo* animal protection studies. W.H., R.M., G.S., K.D., L.V.T., D.R.M., A.S., S.C., P.Y., N.B., J.L.T., R.M.V., P.Z., M.Y. H.L., F.A., M.P., E.G., I.A.W., A.B.W., T.F.R., R.S.B., L.E.G., D.R.B. and R.A. designed the experiments and/or analyzed the data. W.H., R.M., D.R.B. and R.A. wrote the paper, and all authors reviewed and edited the paper.

## Methods

### Convalescent COVID-19 and human vaccinee sera

Sera from convalescent COVID-19 donors ^29^, spike-mRNA-vaccinated humans, and from COVID-19-recovered vaccinated donors, were provided through the “Collection of Biospecimens from Persons Under Investigation for 2019-Novel Coronavirus Infection to Understand Viral Shedding and Immune Response Study” UCSD IRB# 200236. The protocol was approved by the UCSD Human Research Protection Program. Convalescent serum samples were collected based on COVID-19 diagnosis regardless of gender, race, ethnicity, disease severity, or other medical conditions. All human donors were assessed for medical decision-making capacity using a standardized, approved assessment, and voluntarily gave informed consent prior to being enrolled in the study. The summary of the demographic information of the COVID-19 convalescent and vaccinated donors is listed in Table S1.

### Plasmid construction

To generate soluble S ectodomain proteins from SARS-CoV-1 (residues 1-1190; GenBank: AAP13567) and SARS-CoV-2 (residues 1-1208; GenBank: MN908947), we constructed the expression plasmids by synthesizing the DNA fragments from GeneArt (Life Technologies) and cloned them into the phCMV3 vector (Genlantis, USA). To keep the soluble S proteins in a stable trimeric prefusion state, the following changes in the constructs were made: double proline substitutions (2P) were introduced in the S2 subunit; the furin cleavage sites (in SARS-CoV-2 residues 682–685, and in SARS-CoV-1 residues 664–667) were replaced by “GSAS” linker; the trimerization motif T4 fibritin was incorporated at the C-terminus of the S proteins. To purify and biotinylate the spike proteins, the HRV-3C protease cleavage site, 6x HisTag, and AviTag spaced by GS-linkers were added to the C-terminus after the trimerization motif. To produce truncated proteins of SARS-CoV-1 and SARS-CoV-2 spike, the PCR amplifications of the gene fragments encoding SARS-CoV-1 RBD (residue 307-513), SARS-CoV-2 NTD (residue 1-290), RBD (residue 320-527), RBD-SD1 (residue 320-591), and RBD-SD1-2 (residue 320-681) subdomains were carried out using the SARS-CoV-1 and SARS-CoV-2 plasmids as templates. To generate pseudoviruses of non-human sarbecoviruses, the DNA fragments encoding the spikes of the sarbecoviruses without the ER retrieval signal were codon-optimized and synthesized at GeneArt (Life Technologies). The spike encoding genes of Pang17 (residues 1-1249, GenBank: QIA48632.1), WIV1 (residues 1-1238, GenBank: KF367457) and SHC014 (residue 1-1238, GenBank: AGZ48806.1) were constructed into the phCMV3 vector (Genlantis, USA) using the Gibson assembly (NEB, E2621L) according to the manufacturer’s instructions. To express the monomeric RBDs of sarbecovirus clades (clades, 1b, 1a, 2 and 3), the conserved region aligning to SARS-CoV-2 RBD (residue 320-527) were constructed into phCMV3 vector with 6x HisTag, and AviTag spaced by GS-linkers on C-terminus. The sarbecovirus RBD genes encoding RaTG13 (residues 320-527, GenBank: QHR63300.2), Pang17 (residues 318-525, GenBank: QIA48632.1), WIV1 (residues 308-514, GenBank: KF367457), RsSHC014 (residues 308-514, GenBank: AGZ48806.1), BM-4831 (residues 311-514, NCBI Reference Sequence: NC_014470.1), BtKY72 (residues 310-516, GenBank: KY352407), RmYN02 (residues 299-487, GSAID EPI_ISL_412977), Rf1 (residues 311-499, GenBank: DQ412042.1), Rs4081 (residues 311-499, GenBank: KY417143.1) and Yun11 (residues 311-499, GenBank: JX993988) were synthesized at GeneArt (Life Technologies) and constructed using the Gibson assembly (NEB, E2621L).

### Cell lines

HEK293F cells (Life Technologies) and Expi293F cells (Life Technologies) were maintained using 293FreeStyle expression medium (Life Technologies) and Expi293 Expression Medium (Life Technologies), respectively. HEK293F and Expi293F cell suspensions were maintained in a shaker at 150 rpm, 37°C with 8% CO_2_. Adherent HEK293T cells were grown in DMEM supplemented with 10% FBS and 1% penicillin-streptomycin and maintained in an incubator at 37°C with 8% CO_2_. A stable hACE2-expressing HeLa cell line was generated using an ACE2 lentivirus protocol previously described. Briefly, the pBOB-hACE2 plasmid and lentiviral packaging plasmids (pMDL, pREV, and pVSV-G (Addgene #12251, #12253, #8454)) were co-transfected into HEK293T cells using the Lipofectamine 2000 reagent (ThermoFisher Scientific, 11668019).

### Transfection for protein expression

For expression of mAbs, HC and LC gene segments that were cloned into corresponding expression vectors were transfected into Expi293 cells (Life Technologies) (2-3 million cells/mL) using FectoPRO PolyPlus reagent (Polyplus Cat # 116-040) for a final expression volume of 2, 4 or 50 mL. After approximately 24 hours, sodium valproic acid and glucose were added to the cells at a final concentration of 300 mM each. Cells were allowed to incubate for an additional 4 days to allow for mAb expression. For expression of spike proteins, RBDs, and NTDs, cloned plasmids (350 μg) were transfected into HEK293F cells (Life Technologies) (1 million cells/mL) using Transfectagro reagent (Corning) and 40K PEI (1 mg/mL) in a final expression volume of 1 L as previously described. Briefly, plasmid and transfection reagents were combined and filtered preceding PEI addition. The combined transfection solution was allowed to incubate at room temperature for 30 mins before being gently added to cells. After 5 days, supernatant was centrifuged and filtered.

### Protein purification

For mAb purification, a 1:1 solution of Protein A Sepharose (GE Healthcare) and Protein G Sepharose (GE Healthcare) was added to Expi293 supernatant for 2h at room temperature or overnight at 4°C. The solution was then loaded into an Econo-Pac column (BioRad #7321010), washed with 1 column volume of PBS, and mAbs were eluted with 0.2 M citric acid (pH 2.67). The elution was collected into a tube containing 2 M Tris Base. Buffer was exchanged with PBS using 30K Amicon centrifugal filters (Millipore, UFC903008). His-tagged proteins were purified using HisPur Ni-NTA Resin (Thermo Fisher). Resin-bound proteins were washed (25 mM Imidazole, pH 7.4) and slowly eluted (250 mM Imidazole, pH 7.4) with 25 mL elution buffer. Eluted proteins were buffer-exchanged with PBS, and further purified using size-exclusion chromatography using Superdex 200 (GE Healthcare).

### ELISA

ELISAs were performed on 96-well half-area microplates (ThermoFisher Scientific) as described previously ^15^. The plate was coated with 2 µg/mL mouse anti-His antibody (Invitrogen cat. #MA1-21315-1MG, ThermoFisher Scientific) overnight at 4°C. The following day, plates were washed three times with PBST (PBS + 0.05% Tween20) and incubated for 1h with blocking buffer (3% bovine serum albumin (BSA)). Following removal of blocking buffer, plates were treated with His-tagged proteins (5 µg/mL in PBST + 1% BSA) for 1.5h at room temperature. Plates were washed and serum was added at threefold dilutions (beginning at 1:30) and allowed to incubate for 1.5h. Following washes, secondary antibody (AffiniPure Goat anti-human IgG Fc fragment specific, Jackson ImmunoResearch Laboratories cat. #109-055-008) was added for an additional 1h. Secondary antibody was washed, and staining substrate (alkaline phosphatase substrate pNPP tablets, Sigma) was added. Absorbance at 405 nm was measured after 8, 20, and 30 min using VersaMax microplate reader (Molecular Devices) and analyzed using SoftMax version 5.4 (Molecular Devices).

### Biotinylation of proteins

To randomly biotinylate the proteins described in this paper, we used an EZ-Link NHS-PEG Solid-Phase Biotinylation Kit (Thermo Scientific #21440). To dissolve the reagents supplied in the kit for stock solutions, 10 µL DMSO was added into each tube. To make a working solution, 1 µL stock solution was diluted by 170 µL water freshly before use. To concentrate the proteins before biotinylation, the proper sized filter Amicon tubes were used. The proteins were adjusted to 7-9 mg/mL in PBS. For each 30 µL aliquoted protein, 3 µL of working solution was added and mixed thoroughly following by a 3h incubation on ice. To stop the reaction and remove the free NHS-PEG4-Biotin, the protein solution was buffer exchanged into PBS using Amicon tubes. All proteins were evaluated by BioLayer Interferometry after biotinylation.

### BirA biotinylation of proteins for B cell sorting

For B cell sorting, the spike probes with the His and Avi-tag at the C-terminus were biotinylated by the intracellular biotinylating reaction during transfection step. To biotinylate the recombinant Avi-tagged spike probes, the BirA biotin-protein ligase encoding plasmid was co-transfected with the spike probe-Avi-tag encoding plasmids in the FreeStyle™ 293-F cell. 150ug BirA plasmid and 300ug spike probe plasmids were transfected with PEI reagent as described in the Transient transfection section. The spike probes were purified with HisPur Ni-NTA Resin (Thermo Fisher) as described in the Protein purification section. After the purification, the biotinylated proteins were evaluated by BioLayer Interferometry.

### BioLayer Interferometry (BLI)

Binding assays were performed on an Octet RED384 instrument using Anti-Human IgG Fc Capture (AHC) biosensors (ForteBio). All samples were diluted in Octet buffer (PBS with 0.1% Tween 20) for a final concertation of 10 µg/mL for mAbs and 200 nM for viral proteins. For supernatant mAb binding screening, 125 µL of expression supernatant was used. For binding assays, antibodies were captured for 60 s and transferred to buffer for an additional 60 s. Captured antibodies were dipped into viral proteins for 120 s in order to obtain an association signal. For dissociation, biosensors were moved to Octet buffer only for an additional 240 s for the dissociation step. The data generated was analyzed using the ForteBio Data Analysis software for correction, and the kinetic curves were fit to 1:1 binding mode. Note that the IgG: spike protomer binding can be a mixed population of 2:1 and 1:1, such that the term ‘apparent affinity’ dissociation constants (K_D_^App^) are shown to reflect the binding affinity between IgGs and spike trimers tested.

### Isolation of monoclonal antibodies (mAbs)

To isolate antigen-specific memory B cells, we used SARS-CoV-1 and SARS-CoV-2 spike proteins as probes to perform single cell sorting in a 96-well format. PBMCs from post-infection vaccinated human donors were stained with fluorophore labeled antibodies and spike proteins. To generate spike probes, streptavidin-AF647 (Thermo Fisher S32357) was coupled to BirA biotinylated SARS-CoV-1 spike. Streptavidin-AF488 (Thermo Fisher S32354) and streptavidin-BV421 (BD Biosciences 563259) were coupled to BirA biotinylated SARS-CoV-2 spike separately. The conjugation reaction was carried freshly before use with spike protein versus streptavidin-fluorophores at 2:1 or 4:1 molecular ratio. After 30 min incubation at room temperature, the conjugated spike proteins were stored on ice or at 4 ℃ for up to 1 week. To prepare PBMCs, the frozen PBMCs were thawed in 10mL recover medium (RPMI 1640 medium containing 50% FBS) immediately before staining. The cells were washed with 10mL FACS buffer (PBS, 2% FBS, 2 mM EDTA) and each 10 million cells were resuspended in 100µL of FACS buffer. To isolate SARS-CoV-1 and SARS-CoV-2 cross-reactive IgG+ B cells, PBMCs were stained for CD3 (APC Cy7, BD Pharmingen #557757), CD4 (APC-Cy7, Biolegend, #317418), CD8 (APC-Cy7, BD Pharmingen #557760), CD14 (APC-H7, BD Pharmingen #561384, clone M5E2), CD19 (PerCP-Cy5.5, Biolegend, #302230, clone HIB19), CD20 (PerCP-Cy5.5, Biolegend, #302326, clone 2H7), IgG (BV786, BD Horizon, #564230, Clone G18-145) and IgM (PE, Biolegend, #314508, clone MHM-88). Antibodies were incubated with PBMCs on ice for 15 min. After the 15 min staining, SARS-CoV-1-S-AF647, SARS-CoV-2-S-AF488, and SARS-CoV-2-S-BV421 were added to the PBMC solution incubating on ice. After another 30 min incubation, FVS510 Live/Dead stain (Thermo Fisher Scientific, #L34966) 1:1000 diluted with FACS buffer was added to the PBMC solution for 15 min. Subsequently, cells were washed with 10 mL ice cold FACS buffer. Each 10 million cells were resuspended with 500µL FACS buffer and then filtered through 70um nylon mesh FACS tube caps (Fisher Scientific, #08-771-23). A BD FACSMelody sorter (BRV 9 Color Plate 4way) was used for the single cell sorting process. To isolate cross-reactive B cells, the gating strategy was set as follows: lymphocytes (SSC-A vs. FSC-A) and singlets (FSC-H vs. FSC-A) were gated first, and then live cells were selected by FVS510 Live/Dead negative gating. B cells were identified as CD19+CD20+CD3-CD4-CD8-CD14-IgM-IgG+ live singlets. Cross-reactive S-protein specific B cells were sequentially selected for SARS-CoV-2-S-BV421/SARS-CoV-2-S-AF488 double positivity and SARS-CoV-1-S-AF647/SARS-CoV-2-S-AF488 double positivity. Single cells were sorted into 96-well plates on a cooling platform. To prevent degradation of mRNA, plates were moved onto dry ice immediately after sorting. Reverse transcription was done right after. Superscript IV Reverse Transcriptase (Thermo Fisher), dNTPs (Thermo Fisher), random hexamers (Gene Link), Ig gene-specific primers, DTT, and RNAseOUT (Thermo Fisher), and Igepal (Sigma) were used in the reverse transcription PCR reaction as described previously ^86, 87^. To amplify IgG heavy and light chain variable regions, two rounds of nested PCR reactions were carried out using the cDNAs as template and Hot Start DNA Polymerases (Qiagen, Thermo Fisher) and specific primer sets described previously ^86, 87^. The PCR products of the heavy and light chain variable regions were purified with SPRI beads according to the manufacturer’s instructions (Beckman Coulter). Then, the purified DNA fragments were constructed into expression vectors encoding human IgG1, and Ig kappa/lambda constant domains, respectively. Gibson assembly (NEB, E2621L) was used according to the manufacturer’s instructions in the construction step. To produce mAbs, the paired heavy and light chain were co-transfected into 293Expi cells.

### Immunogenetics analysis

Heavy and light chain sequences of monoclonal antibodies as well as ten Rep-seq libraries representing naive heavy chain repertoires of ten donors (ERR2567178– ERR2567187) from BioProject PRJEB26509 ^85^ were processed using the DiversityAnalyzer tool ^88^. Clonal lineages for mAbs were computed in three steps. The first step was applied to heavy chain sequences following the procedures described previously ^89^. Briefly, heavy chain sequences were combined into the same clonal lineage if (i) they share V and J germline genes, (ii) their CDRH3s have the same lengths, and (iii) their CDRH3s share at least 90% nucleotide identity. At the second step, the same procedure was applied to light chain sequences. Finally, each heavy chain clonal lineage was split according to the clonal lineage assignments of corresponding light chain sequences. Phylogenetic trees derived from heavy chain sequences and IGHD gene segments of mAbs were constructed using the ClusterW2 tool ^90^ and visualized using the Iroki tool ^91^.

### Pseudovirus production

To generate pseudoviruses, plasmids encoding the SARS-CoV-1, SARS-CoV-2 or other variants spike proteins with the ER retrieval signal removed were co-transfected with MLV-gag/pol and MLV-CMV-Luciferase plasmids into HEK293T cells. Lipofectamine 2000 (Thermo Fisher Scientific, 11668019) was used according to the manufacturer’s instructions. 48 hours post transfection, supernatants containing pseudoviruses were collected and filtered through a 0.22 μm membrane to remove debris. Pseudoviruses could be stored at -80°C prior to use.

### Pseudovirus entry and serum neutralization assays

To generate hACE2-expressing stable cell lines for the pseudovirus infection test, we used lentivirus to transduce the hACE2 into HeLa cells. Stable cell lines with consistent and high hACE2 expression levels were established as HeLa-hACE2 and used in the pseudovirus neutralization assay. To calculate the neutralization efficiency of the sera or mAbs, the samples were 3-fold serially diluted and 25 μL of each dilution was incubated with 25 μL of pseudovirus at 37 °C for 1 h in 96-half area well plates (Corning, 3688). Just before the end of the incubation, HeLa-hACE2 cells were suspended with culture medium at a concentration of 2 x 10^5^/mL. The DEAE-dextran (Sigma, # 93556-1G) was added to the cell solutions at 20 μg/mL. 50 μL of the cell solution was distributed into each well. The plates were incubated at 37 °C for 2 days and the neutralization efficiency was calculated by measuring the luciferase levels in the HeLa-hACE2 cells. After removal of the supernatant, the HeLa-hACE2 cells were lysed by luciferase lysis buffer (25 mM Gly-Gly pH 7.8, 15 mM MgSO4, 4 mM EGTA, 1% Triton X-100) at room temperature for 10-20 mins. After adding Bright-Glo (Promega, PRE2620) to each well, luciferase activity was inspected by a luminometer. Each experiment was carried out with duplicate samples and repeated independently at least twice. Percentage of neutralization was calculated according to the equation:

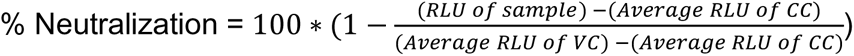

The neutralization percentage was calculated and plots against antibody concentrations or sera dilution ratio were made in Graph Pad Prism. The curves were fitted by nonlinear regression and the 50% pseudovirus neutralizing (IC_50_) or binding (ID_50_) antibody titer was calculated.

### Neutralization assay of replication competent sarbecoviruses

Vero E6 cells (ATCC-C1008) were seeded at 2 x 10^4^ cells/well in a black-well, black-wall, tissue culture treated, 96-well plate (Corning Cat. #3916) 24 h before the assay. MAbs were diluted in MEM supplemented with 5%FBS and 1%Pen/Strep media to obtain an 8-point, 3-fold dilution curve with starting concentration at 20 µg/ml. Eight hundred PFU of SARS1-nLuc, SARS2-D614G-nLuc and SHC014-nLuc replication competent viruses were mixed with mAbs at a 1:1 ratio and incubated at 37°C for 1 h. One-hundred microliters of virus and mAb mix was added to each well and incubated at 37°C + 5% CO_2_ for 20 to 22 h. Luciferase activities were measured by the Nano-Glo Luciferase Assay System (Promega Cat. #N1130) following the manufacturer’s protocol using a GloMax luminometer (Promega). Percent inhibition and IC_50_ were calculated as pseudovirus neutralization assay described above. All experiments were performed as duplicate and independent repeated for three times. All the live virus experiments were performed under biosafety level 3 (BSL-3) conditions at negative pressure, by operators in Tyvek suits wearing personal powered-air purifying respirators.

### Competition BLI

To determine the binding epitopes of the isolated mAbs compared with known human SARS-CoV-2 mAbs, we did in-tandem epitope binning experiments using the Octet RED384 system. 200 nM of randomly biotinylated SARS-CoV-2 S or RBD protein antigen was captured using SA biosensors (18-5019, Sartorius). The biosensor was loaded with antigen for 5 min and then moved into the saturating mAbs at a concentration of 100 µg/mL for 10 min. The biosensors were then moved into bnAb solution for 5 min to measure binding in the presence of saturating antibodies. As control, biosensors loaded with antigen were directly moved into bnAb solution. The percent (%) inhibition in binding is calculated with the formula: [Percent (%) binding inhibition = 1- (bnAb binding response in presence of the competitor antibody / binding response of the corresponding control bnAb without the competitor antibody)]

### Fab production

To generate the Fab from the IgG, a stop codon was inserted in the heavy chain constant region at “KSCDK”. The truncated heavy chains were co-transfected with the corresponding light chains in 293Expi cells to produce the Fabs. The supernatants were harvested 4 days post transfection. Fabs were purified with CaptureSelect™ CH1-XL MiniChrom Columns (#5943462005). Supernatants were loaded onto columns using an Econo Gradient Pump (Bio-Rad #7319001). Following a wash with 1x PBS, Fabs were eluted with 25 mL of 50 mM acetate (pH 4.0) and neutralized with 2 M Tris Base. The eluate was buffer exchanged with 1x PBS in 10K Amicon tubes (Millipore, UFC901008) and filtered with a 0.22 µm spin filter.

### Negative stain electron microscopy

S-protein was complexed with Fab at three times molar excess per trimer and incubated at room temperature for 30 mins. Complexes were diluted to 0.03mg/ml in 1x Tris-buffered saline and 3µl applied to a 400mesh Cu grid, blotted with filter paper, and stained with 2% uranyl formate. Micrographs were collected on a Thermo Fisher Tecnai Spirit microscope operating at 120kV with an FEI Eagle CCD (4k x 4k) camera at 52,000 X magnification using Leginon automated image collection software ^92^. Particles were picked using DogPicker ^93^ and data was processed using Relion 3.0 ^94^. Map segmentation was performed in UCSF Chimera ^95^.

### In vivo infections

All mouse experiments were performed at the University of North Carolina, NIH/PHS Animal Welfare Assurance Number: D16-00256 (A3410-01), under approved IACUC protocols. All animal manipulation and virus work was performed in a Class 2A biological safety cabinet. 12 month old female Balb/c mice (strain 047) were purchased from Envigo. Mice were housed in individually ventilated Seal-Safe cages, provided food and water ad libitum and allowed to acclimate at least seven days before experimental use. Twelve hours prior to infection, mice were intraperitoneally injected with 300μg of antibody. Immediately prior to infection, mice were anesthetized by intraperitoneal injection of ketamine and xylazine and weighed. Virus was diluted in 50μL of sterile PBS and administered intranasally. Mice were weighed daily and observed for signs of disease. At the designated timepoint, mice were euthanized via isoflurane overdose, gross lung pathology was assessed, and the inferior lobe was collected for virus titration. Respiratory function was measured at day two post infection via Buxco whole body plethysmography, as previously described ^96^.

### Virus titration

SARS-CoV-2 MA10, SARS-CoV-1 MA15 and chimeric SARS-CoV-1 MA15-SHC014 were grown and titered using VeroE6 cells as previously described ^97^. Briefly, lung tissue was homogenized in 1mL sterile PBS via Magnalyser (Roche), centrifuged to pellet debris, plated in 10-fold serial dilutions on VeroE6 cells on a 6-well plate and covered with a 1:1 mixture of 1.6% agarose and media. At two (SARS-CoV-1) or three (SARS-CoV-2) days post plating, cells were stained with neutral red and plaques counted.

### Statistical Analysis

Statistical analysis was performed using Graph Pad Prism 8 for Mac, Graph Pad Software, San Diego, California, USA. Statistical comparisons between the two groups were performed using a Mann-Whitney two-tailed test. The correlation between two groups was determined by Spearman rank test. Groups of data were compared using the Kruskal-Wallis non-parametric test. Dunnett’s multiple comparisons test were also performed between experimental groups. Data were considered statistically significant at * p < 0.05, ** p < 0.01, *** p < 0.001, and **** p < 0.0001.

### Data availability

The authors declare that the data supporting the findings of this study are available within the paper and its supplementary information files or from the corresponding author upon reasonable request. Antibody sequences have been deposited in GenBank under accession numbers OM467906 - OM468119. Antibody plasmids are available from Raiees Andrabi or Dennis Burton under an MTA from The Scripps Research Institute.

---

**Table S1.**
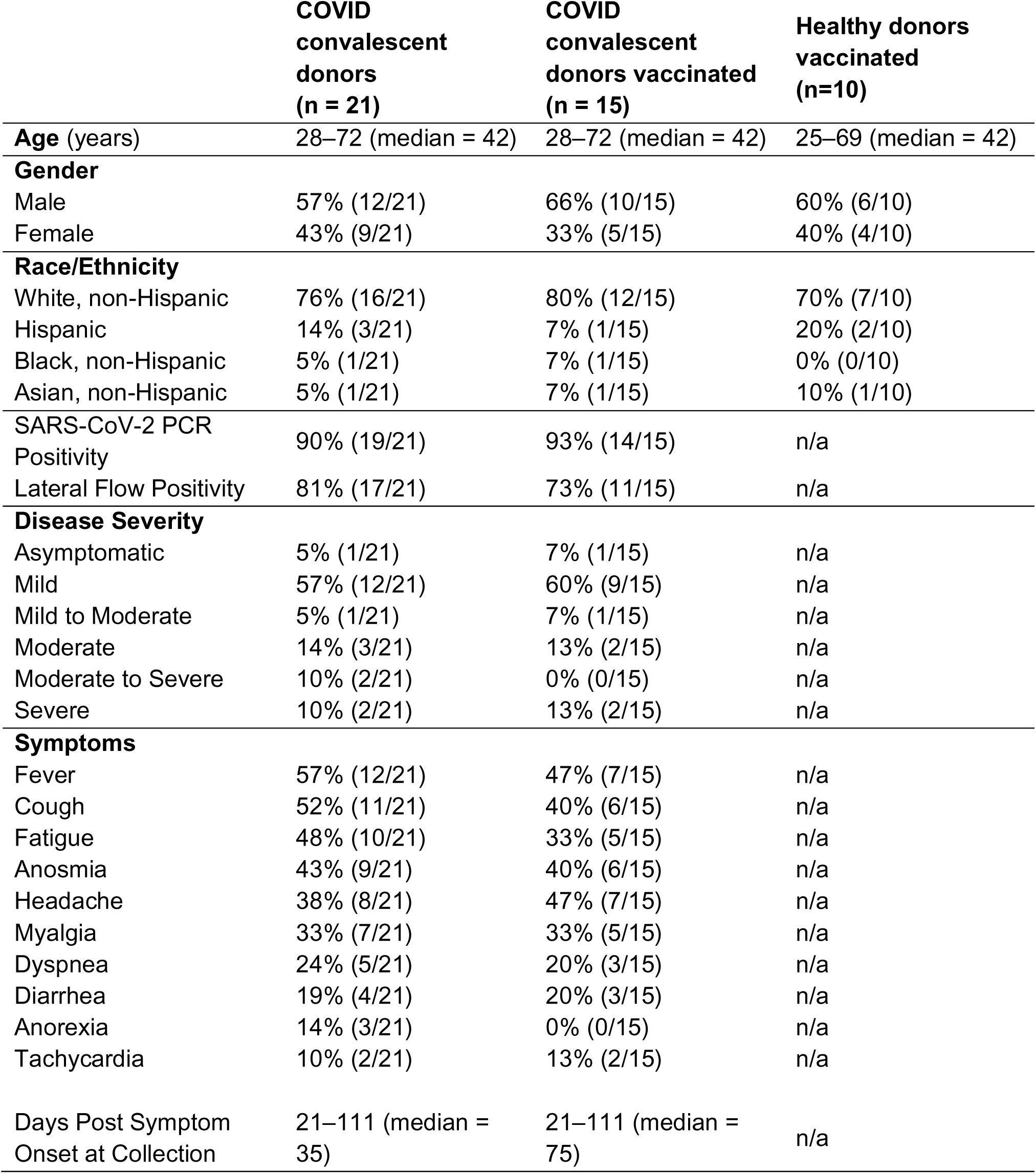
**Demographic information of human donors**

**Extended Data Fig. 1.**
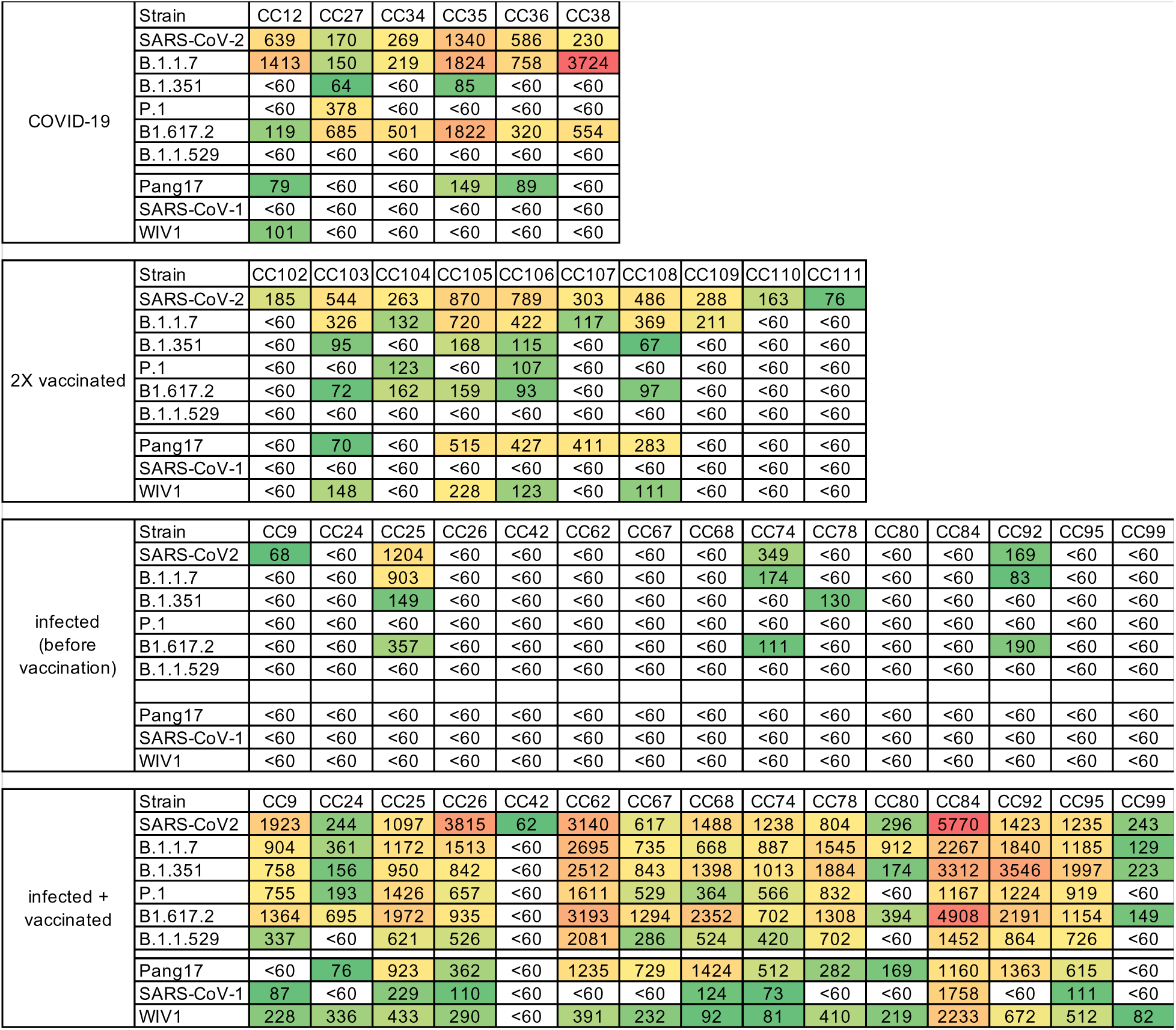
Serum neutralization. Neutralization by sera from COVID-19, 2 x mRNA-spike-vaccinated and SARS-CoV-2 recovered/mRNA vaccinated donors with pseudotyped SARS-CoV-2, SARS-CoV-2 variants of concern [B.1.1.7 (Alpha), B.1.351 (Beta), P.1 (Gamma), B.1.617.2 (Delta) and B.1.1.529 (Omicron)], as well as other sarbecoviruses (Pang17, SARS-CoV-1, and WIV1). ID50 neutralization titers are shown. Prior to vaccination, the sera from infected-vaccinated donors were tested for neutralization and the ID50 neutralization titers are shown for comparison.

**Extended Data Fig. 2.**
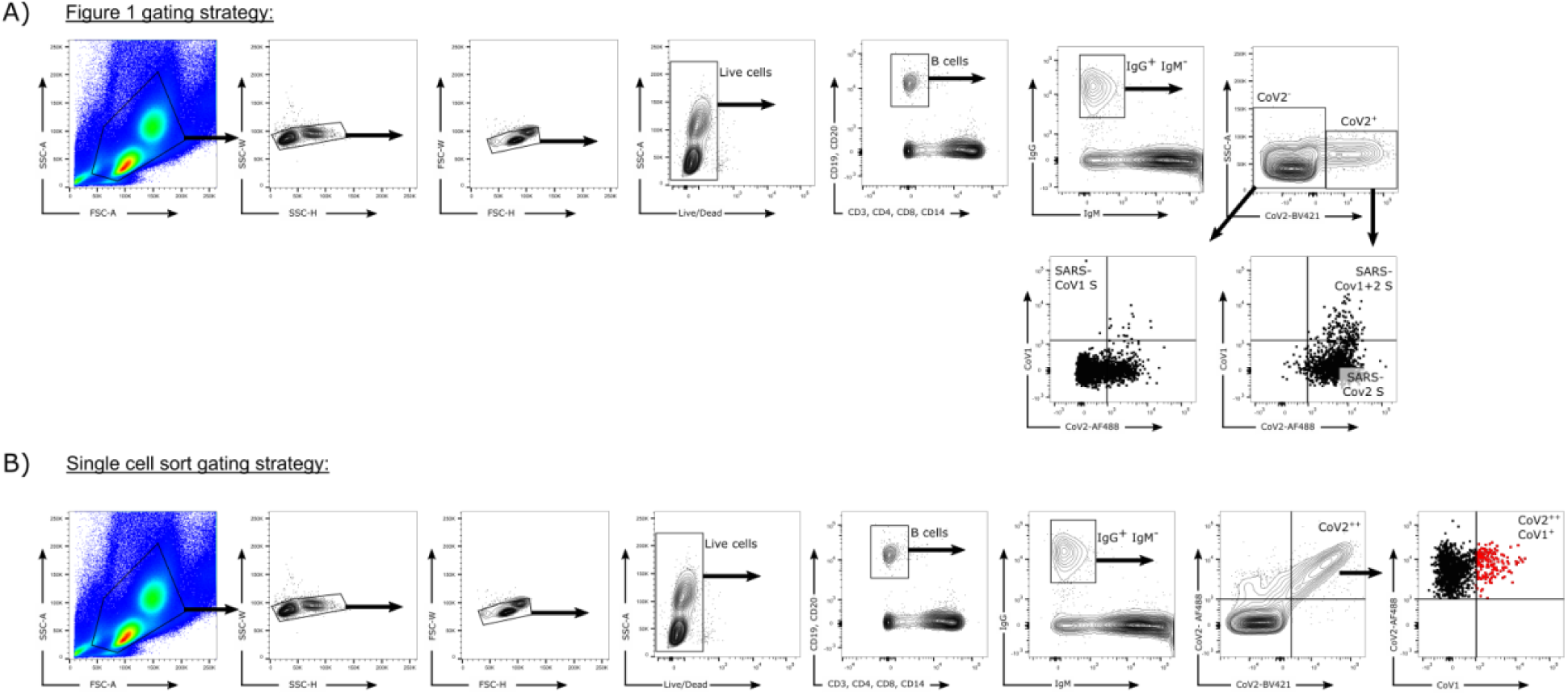
Flow cytometry B cell profiling and sorting strategies. **a.** Gating strategy for analysis of IgG^+^ B cell populations that bind SARS-CoV-1 S-protein only (CD19^+^CD20^+^CD3^-^CD4^-^CD8^-^CD14^-^IgM^-^IgG^+^CoV2BV421^-^CoV2AF488^-^CoV1^+^), SARS-CoV-2 S-protein only (CD19^+^CD20^+^CD3^-^CD4^-^CD8^-^CD14^-^IgM^-^IgG^+^ CoV2BV421^+^CoV2AF488^+^CoV1^-^), or both SARS-CoV-1 and SARS-CoV-2 S-proteins (CD19^+^CD20^+^CD3^-^CD4^-^CD8^-^CD14^-^IgM^-^ IgG^+^CoV2BV421^+^CoV2AF488^+^CoV1^+^). **b.** Gating strategy used to isolate single cross-reactive IgG^+^ B cells (indicated in red).

**Extended Data Fig. 3.**
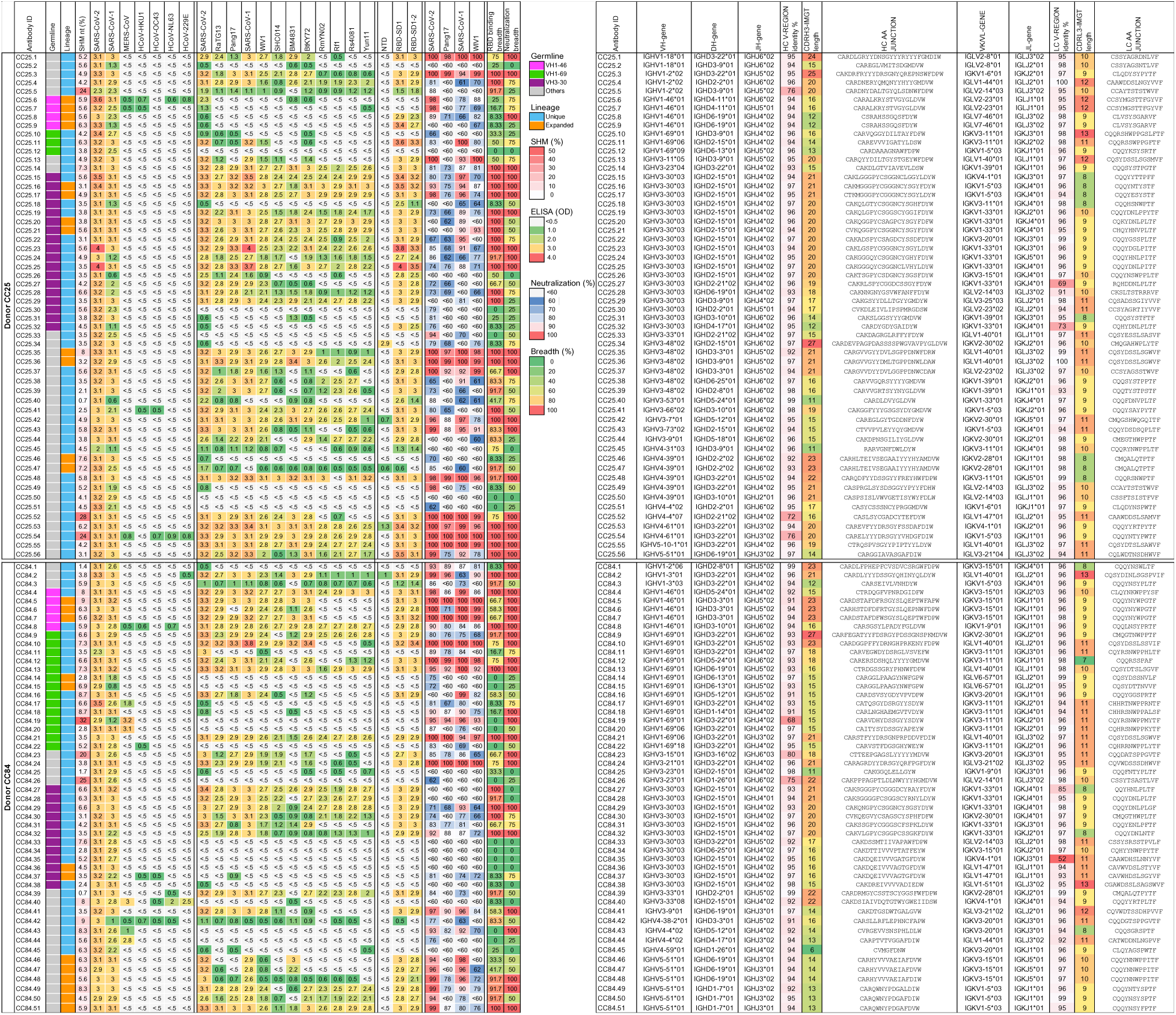
Binding, neutralization and immunogenetics information of isolated mAbs. A total of 107 mAbs from two SARS-CoV-2 recovered-vaccinated donors CC25 (n = 56 mAbs) and CC84 (n = 51 mAbs) were isolated by single B cell sorting using SARS-CoV-1 and SARS-CoV-2 S-proteins as baits. MAbs were expressed and tested for antigen binding, pseudovirus neutralization, and analyzed for immunogenetic properties. Germline, lineage, somatic hypermutation (SHM), ELISA binding to S-proteins and RBDs, neutralization of ACE2-utilizing sarbecoviruses and breadth are colored according to the key. Paired gene information, including heavy chain CDRH3 and light chain CDRL3 sequences are represented for each mAb.

**Extended Data Fig. 4.**
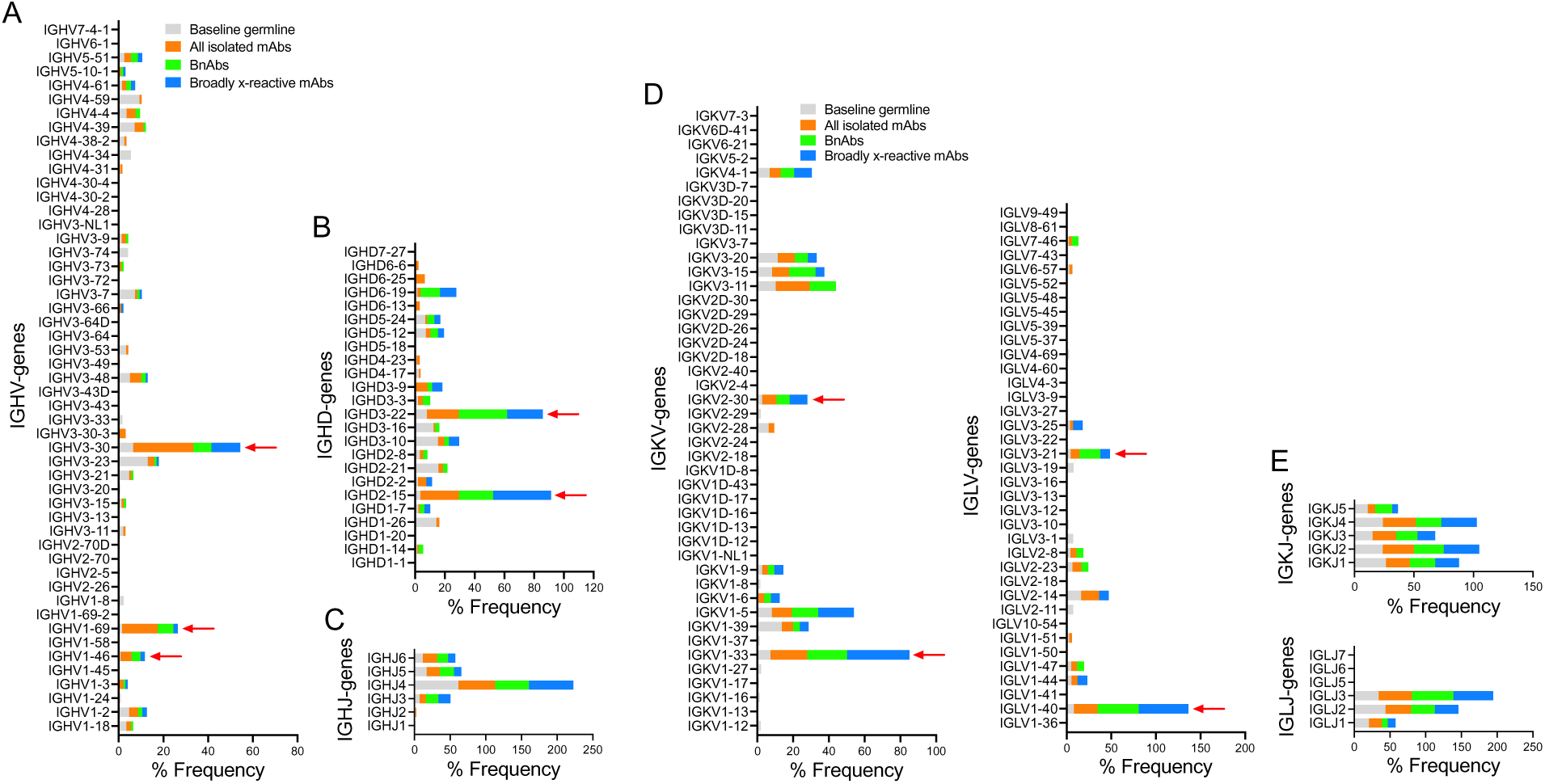
Immunoglobulin heavy and light chain germline gene enrichments in isolated RBD mAbs compared to a reference human germline database. Baseline germline frequencies of heavy chain genes (IGHV (a), IGHD (b) and IGHJ (c) genes) and light chain genes (IGKV and IGLV (d), IGKJ and IGLJ (e) genes) are shown in grey, and mAb, bnAbs and cross-reactive mAbs in a-e panels are colored according to the key in (a and d). Arrows indicate gene enrichments compared to human baseline germline frequencies. The gene usage enrichments in panels a-e are shown for all unique clone mAbs isolated from CC25 and CC84 donors.

**Extended Data Fig. 5.**
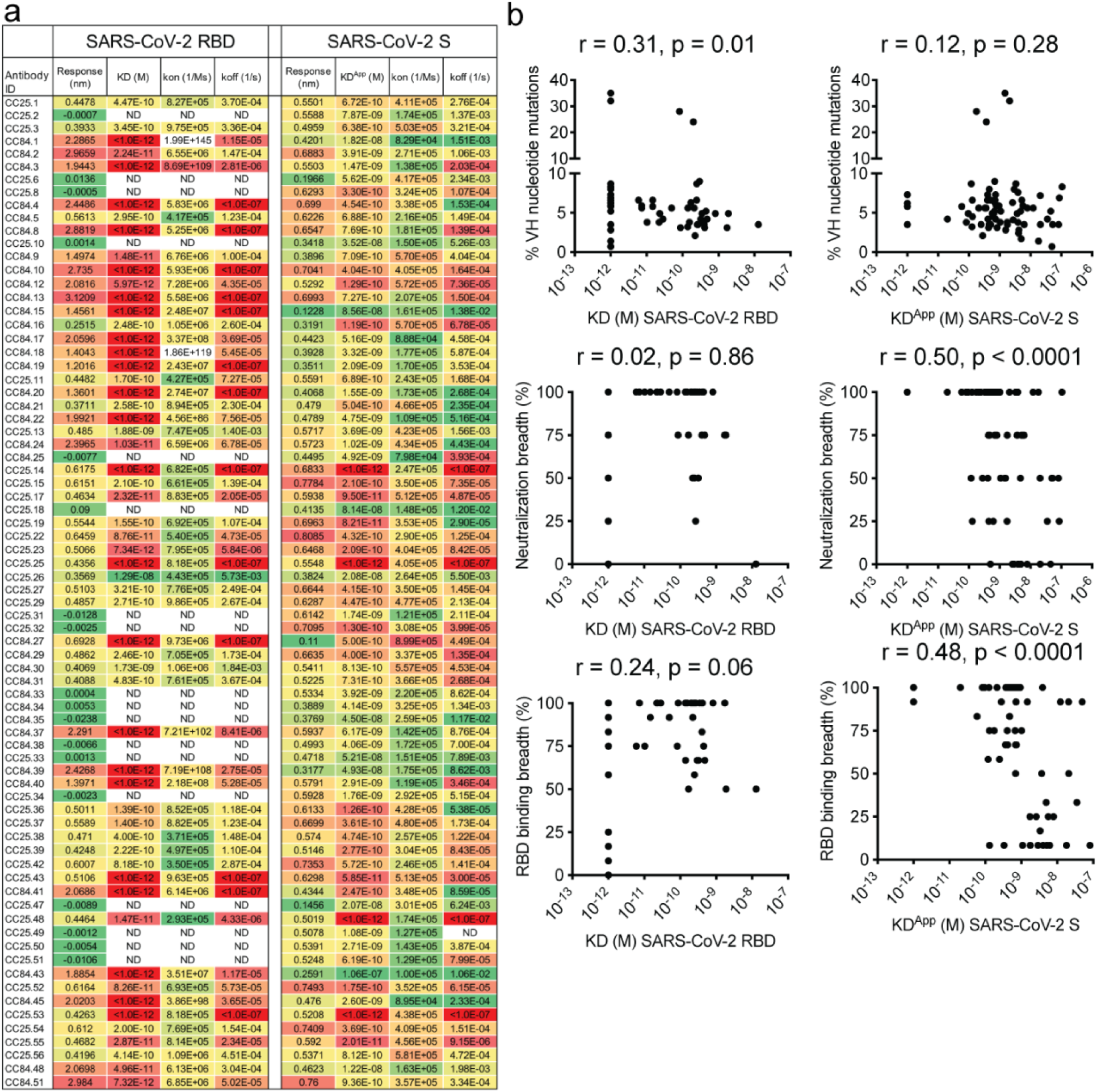
mAb supernatant binding to SARS-CoV-2 RBD and SARS-CoV-2 S and association with SHM, binding and neutralization breadth. **a.** Supernatants from Expi293F cell-expressed mAbs were screened for BLI binding with SARS-CoV-2 RBD and SARS-CoV-2 S-protein. Binding kinetics (KD (monomeric SARS-CoV-2 RBD) KD^App^ (SARS-CoV-2 S-protein), *k_on_* and *k_off_* constants) of antibodies with human proteins are shown. Binding kinetics were obtained using the 1:1 binding kinetics fitting model on ForteBio Data Analysis software. **b.** Correlations of mAb binding (*K*_D_ or K_D_^App^ (M) values) to SARS-CoV-2 RBD or S-protein with heavy chain SHMs, neutralization breadth (neutralization against 4 ACE2-using-sarbecovirus panel), and sarbecovirus RBD breadth (binding against all 12 RBDs of clades 1a, 1b, 2 and 3) are determined by nonparametric Spearman correlation two-tailed test with 95% confidence interval. The Spearman correlation coefficient (r) and p-value are indicated.

**Extended Data Fig. 6.**
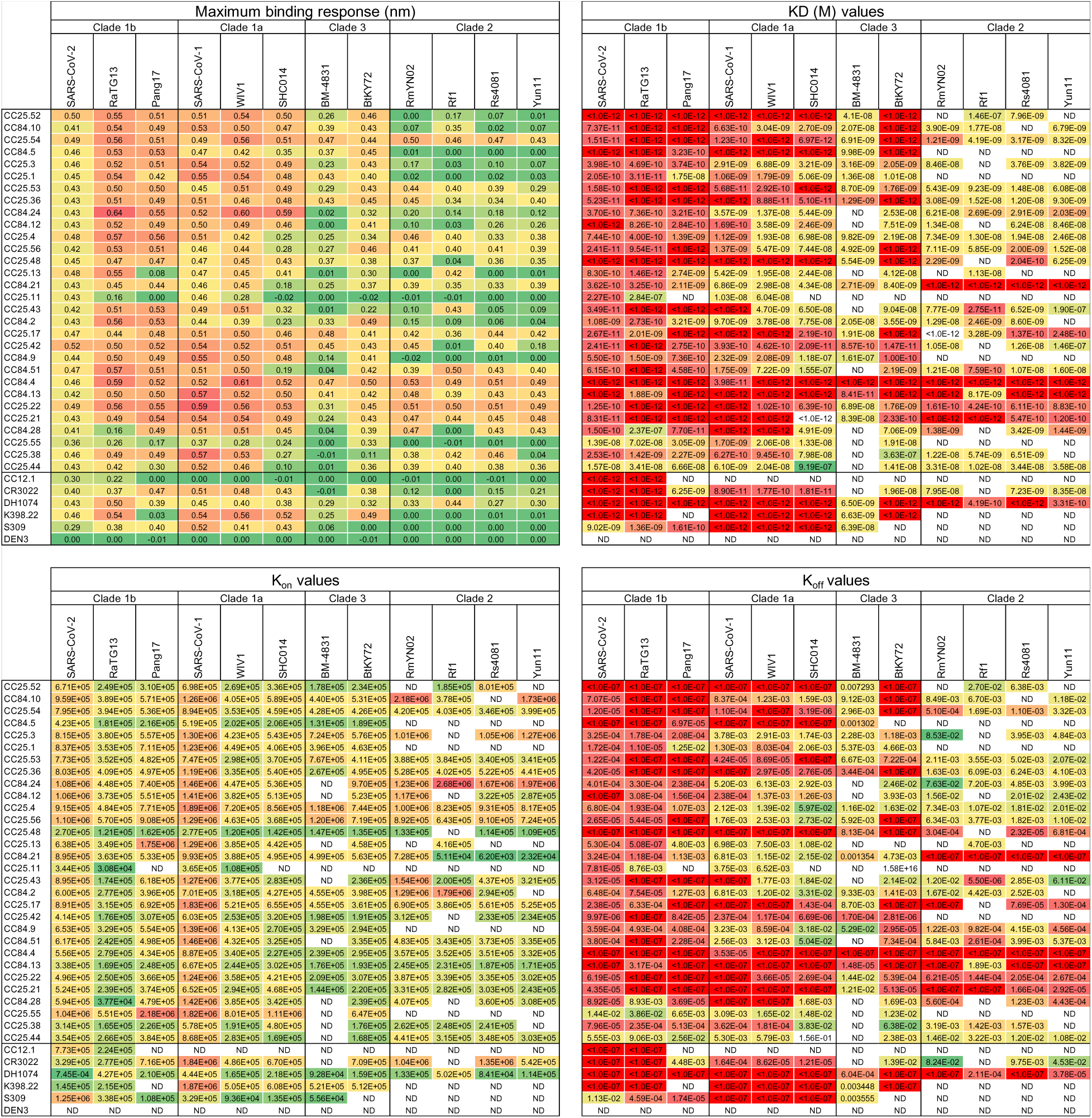
Binding of select mAbs to RBDs from sarbecovirus clades. BLI binding kinetics of select CC25 and CC84 mAbs to monomeric RBDs derived from sarbecovirus clades: clade 1b (SARS-CoV-2, RatG13, Pang17), clade 1a (SARS-CoV-2, WIV1, SHC014), clade 3 (BM-4831, BtKY72) and clade 2 ((RmYN02, Rf1, Rs4081, Yun11). Binding kinetics were obtained using the 1:1 binding kinetics fitting model on ForteBio Data Analysis software and maximum binding responses, dissociations constants (*K*_D_) and on-rate (*k_on_*) and off-rate constants (*k_off_*) for each antibody protein interaction are shown. KD, kon and koff values were calculated only for antibody-antigen interactions where a maximum binding response of 0.05nm was obtained.

**Extended Data Fig. 7.**
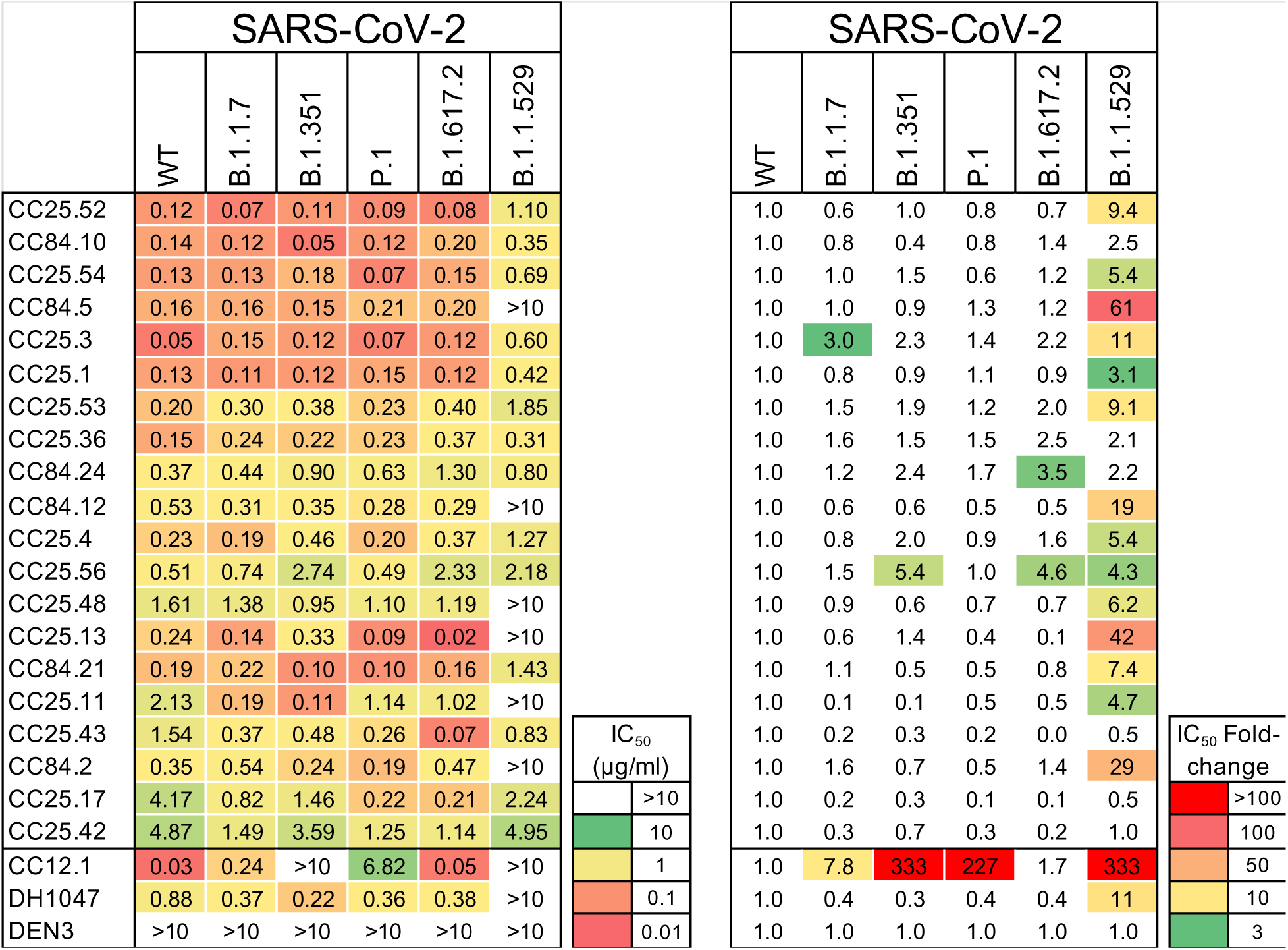
Neutralization of RBD bnAbs against SARS-CoV-2 and major variants of concern (VOCs). IC_50_ neutralization titers of RBD bnAbs against SARS-CoV-2 (WT) and five major SARS-CoV-2 VOCs: (B.1.1.7 (Alpha), B.1.351 (Beta), P.1 (Gamma), B.1.617.2 (Delta) and B.1.1.529 (Omicron)). The IC_50_ neutralization fold-change of RBD bnAbs with SARS-CoV-2 VOCs compared to the WT virus. CC12.1, DH1047 and DEN mAbs were used as controls.

**Extended Data Fig. 8.**
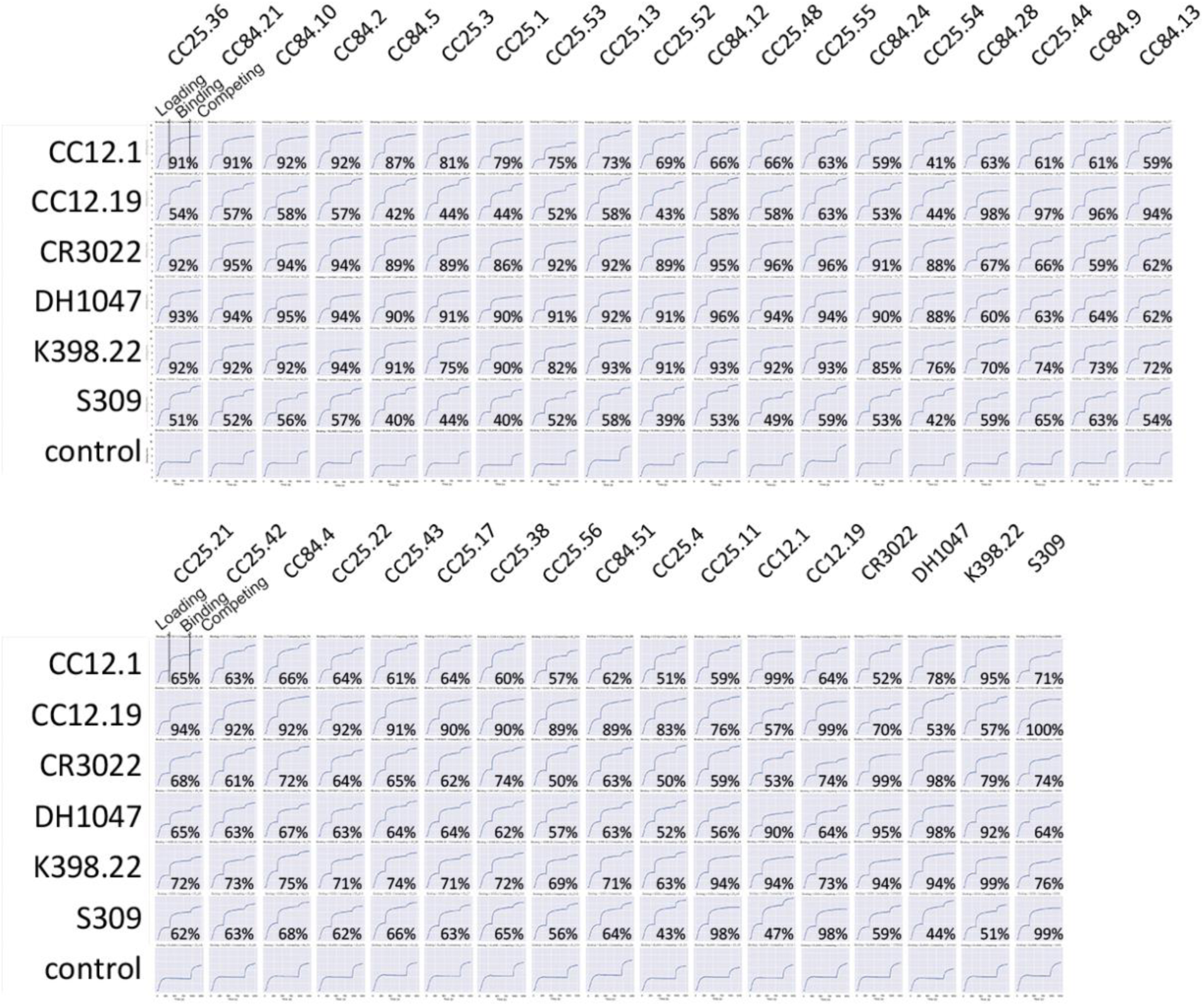
Epitope binning of mAbs using a competition assay. 30 select mAbs (19 mAbs from donor CC25 and 11 mAbs from donor CC84) were assayed in BLI competition binning to evaluate epitope properties shared with previously isolated human (CC12.1, CC12.19, CR3022, DH1047 and S309) and macaque (K398.22) mAbs with known epitope specificities. His-tagged SARS-CoV-2 RBD protein (200nM) was captured on anti-His biosensors and incubated with the indicated mAbs at a saturating concentration of 100µg/mL for 10 mins and followed by nAb incubation for 5 min at a concentration of 25µg/mL. All BLI measurements were performed on an Octet RED384 system. BLI traces are shown for each binding. The binding inhibition % is calculated with the formula: percent (%) of inhibition in the BLI binding response = 1- (response in presence of the competitor antibody / response of the corresponding control antibody without the competitor antibody).

**Extended Data Fig. 9.**
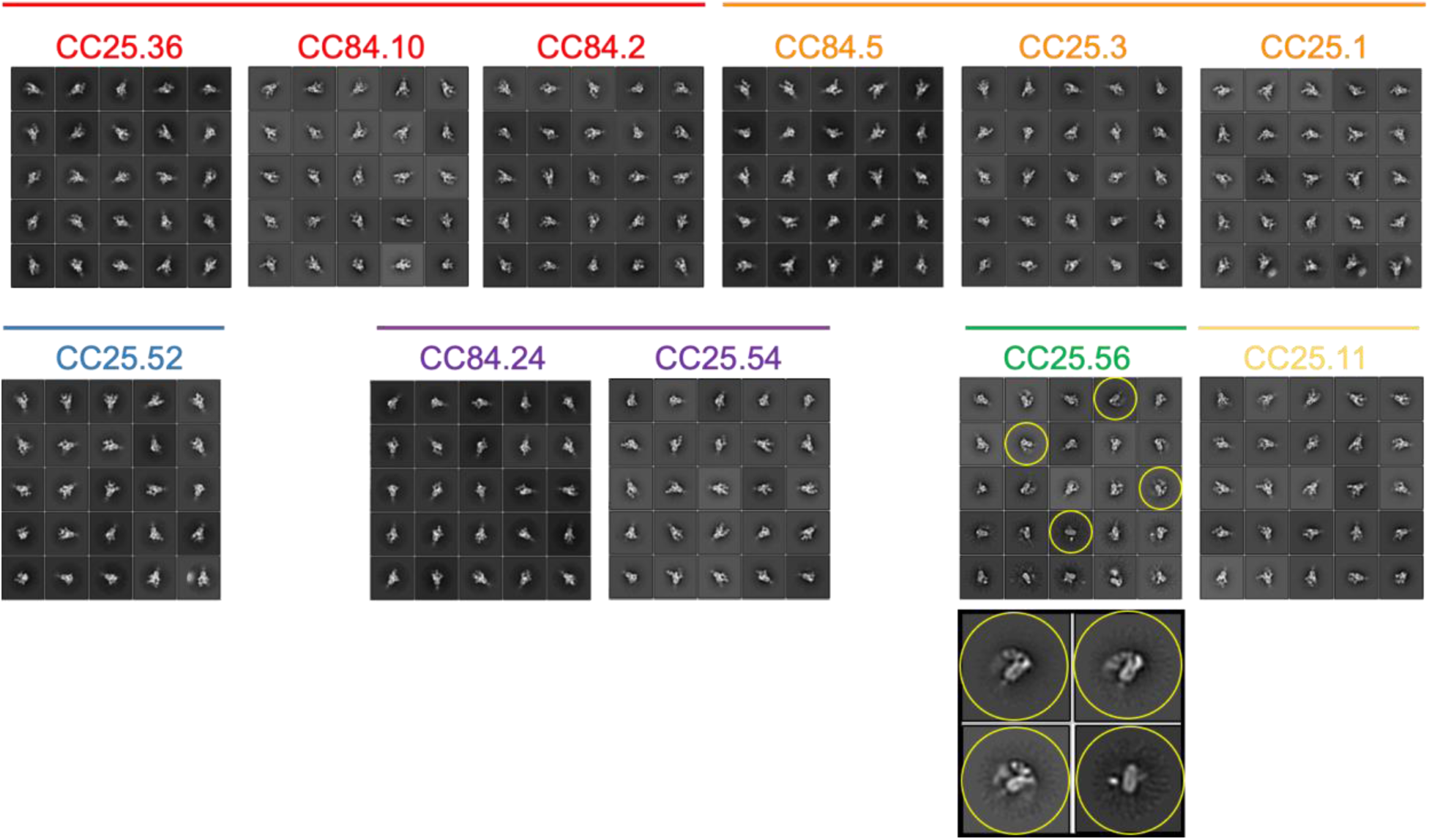
Epitope mapping of bnAbs using negative stain Electron Microscopy (ns-EM). Electron microscopy (EM) images of sarbecovirus cross-neutralizing antibody Fabs with SARS-CoV-2 S-protein. 2D class averages of S-protein bound Fabs for each mAbs are shown. One of the group 2 bnAb Fabs, CC25.56, had some destabilizing effect on the S-protein trimer (indicated in yellow circles).

**Extended Data Fig. 10.**
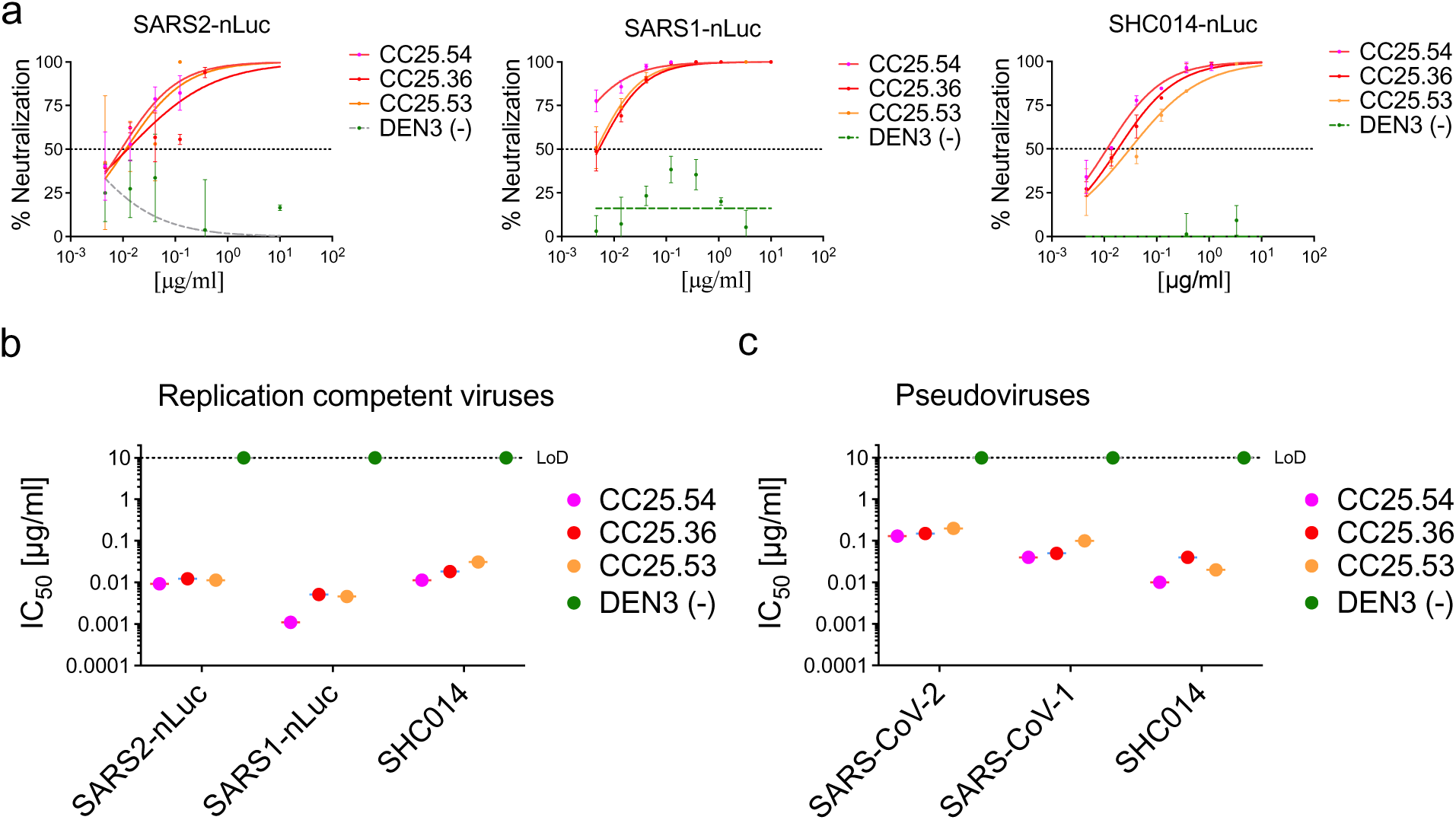
Neutralization of replication competent sarbecoviruses by select RBD bnAbs. **a.** Neutralization of replication competent viruses encoding SARS-CoV-2 (SARS2-nLuc), SARS-CoV-1 (SARS1-nLuc) and SHC014 (SHC014-nLuc) by 3 select RBD bnAbs, CC25.54, CC25.36 and CC25.53. DEN3 antibody was a negative control for the neutralization assay. **b-c**. Comparison of IC_50_ neutralization titers of 3 RBD bnAbs with replication-competent (**b**) and pseudoviruses (**c**) of SARS-CoV-2, SARS-CoV-1 and SHC014 ACE2-utilizing sarbecoviruses.

